# Spindle size scaling secures mitotic fidelity against whole-genome doubling-associated mechanical stress

**DOI:** 10.64898/2026.10.09.757450

**Authors:** Qijiao Wang, Chiharu Watanabe, Sumire Ishida-Ishihara, Yukako Nishimura, Takahiro Yamamoto, Fumio Motegi, Ryota Uehara

## Abstract

Although whole-genome doubling (WGD) accompanies a drastic increase in chromosome number, post-WGD cells often continue to divide. How mitosis is sustained under increased chromosome number remains largely unknown. Here, we found that WGD cells require longer mitotic spindles than isogenic diploids to maintain spindle bipolarity. WGD increased mechanical stress on spindle poles during metaphase, associated with a higher risk of pole fragmentation. Disrupting kinetochore-microtubule attachment alleviated pole stress and fragmentation in WGD cells, suggesting that WGD-linked mechanical stress arises from spindle-chromosome interactions. Artificial spindle elongation reduced the pole stress and mitigated pole fragmentation defects in WGD cells. Our findings identify chromosome gain as a critical mechanical challenge to spindle organization and establish spindle size scaling as an adaptive mechanism that preserves mitotic fidelity after WGD.

## Background

WGD occurs when a cell fails to divide after DNA replication during the cell cycle, resulting in a doubled chromosome number. Many post-WGD cells continue to proliferate and generate diverse progeny, contributing to biological processes such as tumorigenesis and speciation (*1–4*). From the perspective of cell division control, WGD is an irregular event that compromises the accuracy of the process. During normal cell division, two centrosomes form a bipolar spindle, which mediates accurate chromosome congression and segregation. After de novo WGD, doubling the centrosome number (from 2 to 4) often leads to multipolar or unequal chromosome segregation (*5, 6*), which induces chromosome instability. Excess centrosomes are gradually lost during subsequent proliferation (*7, 8*). However, WGD cells with normal centrosome number still show frequent spindle defects and chromosome missegregation (*9–12*), suggesting that increased chromosome number per se has detrimental effects on mitotic control. Despite these observations, it remains largely unclear how increased chromosome number affects mitotic processes and how WGD cells sustain cell division with increased chromosomes. Better understanding of mitotic control after WGD could inform strategies to selectively target tumor-promoting WGD cells.

WGD generally increases cellular content and cell size by just under twofold (*13–16*), causing size scaling of various intracellular structures (*17*). During mitosis, WGD cells often form larger spindles than isogenic diploids (*18–21*). Spindle size scaling has been extensively studied in developmental processes, where spindle sizes adapt to rapid reductions in cell size to ensure robust chromosome segregation through successive early cell divisions (*22–29*). In contrast, whether WGD-linked spindle enlargement has functional significance for mitotic regulation or merely represents a passive epiphenomenon remains unknown.

## Results

### WGD increases TPX2 dependence to maintain mitotic fidelity

We hypothesized that, if any mechanisms secure mitotic fidelity in the WGD state, genes associated with those mechanisms would be more important in WGD cells than in isogenic diploids. Based on this assumption, we compared proliferation between near-diploid human HAP1 cells and their isogenic tetraploid derivatives (*30, 31*), under RNAi-mediated depletion of 52 mitotic genes. We previously generated these tetraploids by pharmacologically inducing WGD via cytokinesis failure, and they lost the excess centrosomes during subsequent clonal isolation (*19*). To enable a direct comparison under identical conditions, we co-cultured GFP-expressing diploids and non-labeled tetraploids at a 1:1 ratio and examined their proportions after 2-day RNAi treatment using flow cytometry (Fig. 1A, S1A, B, and *Supplementary Material S1*; siRNA concentrations were optimized to permit survival of a sufficient number of cells for flow cytometric cell counting).

**Figure 1:**
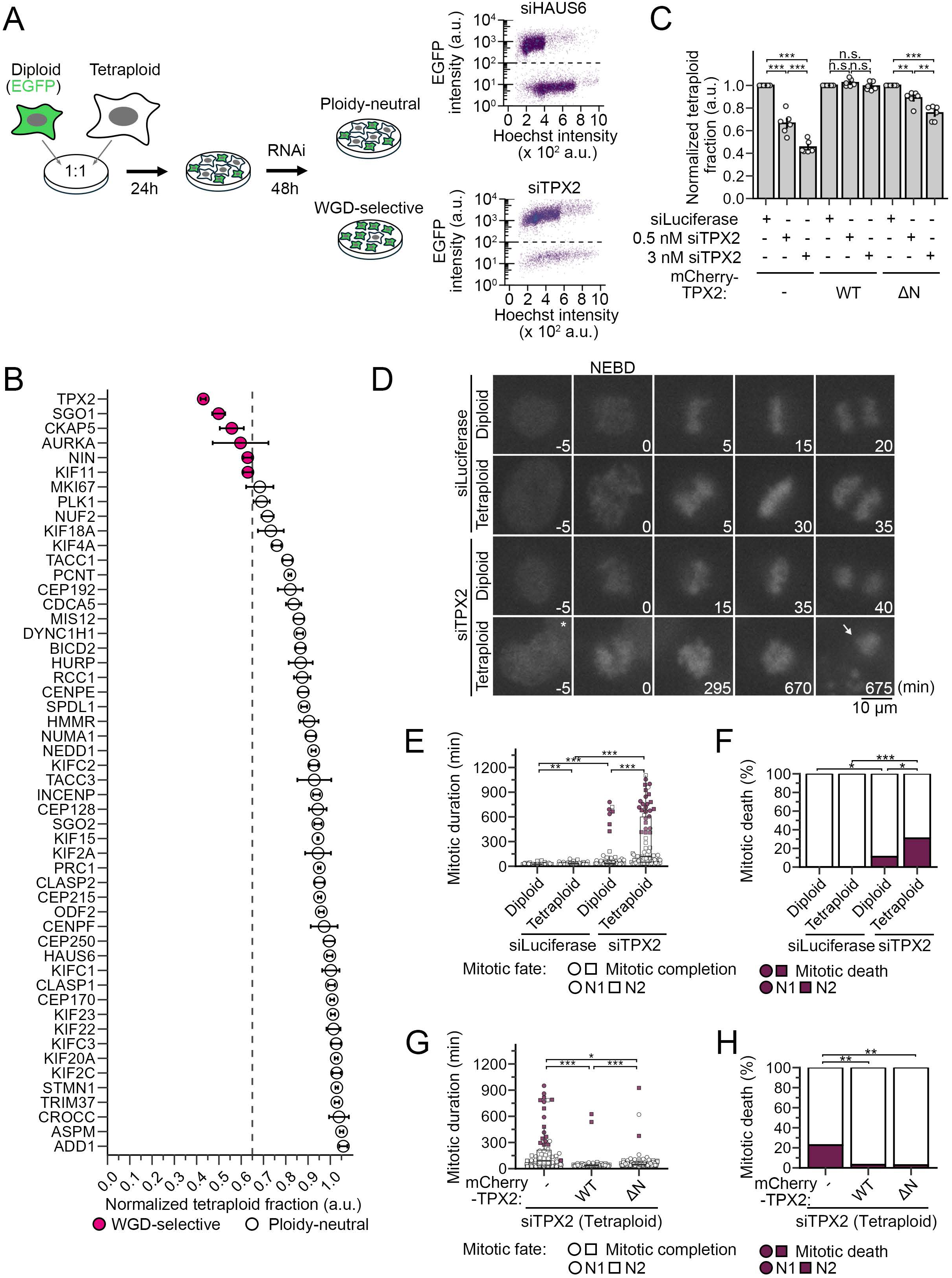
WGD-selective mitotic perturbations by TPX2 depletion in HAP1 cells. **(A)** Left: Scheme of RNAi screen in the 1:1 diploid-tetraploid co-culture. Right: Examples of flow cytometric analysis of EGFP and Hoechst intensity in co-cultures after depletion of a ploidy-neutral (HASU6) or WGD-selective (TPX2) gene. **(B, C)** Normalized tetraploid fraction in the RNAi-treated co-culture relative to the mock-depleted control. Genes with a robust Z-score < 2.5 are categorized “WGD-selective” in B. Indicated RNAi-resistant transgenes were expressed in both diploids and tetraploids in C. Mean ± SE of at least 3 independent experiments. *p<0.05, **p<0.01, ***p<0.001; Tukey–Kramer test for C. **(D)** Time-lapse images of SiR-DNA-stained chromosomes during mitosis in RNAi-treated diploids or tetraploids. Live images were initiated 24 h after siRNA transfection and taken at 5-min intervals. White arrow: mitotic death. Asterisk: a neighboring cell. The timing of NEBD is set as 0 min. **(E-H)** Mitotic duration (from NEBD to mitotic exit; box plots show the median, the 25^th^ and 75^th^ percentiles; E and G) or frequency of mitotic death (F and H). Indicated RNAi-resistant transgenes were expressed both in diploids and tetraploids in G and H. At least 53 cells (E and F) or 66 cells (G and H) pooled from 2 independent experiments were analyzed for each condition. *p<0.05, **p<0.01, ***p<0.001; the DSCF test in E and G, and Fisher exact test in F and H.

Mock-depleted co-culture kept its original ratio of diploids and tetraploids (*Supplementary Material S1*). While depletion of >45 factors had ploidy-neutral effects (i.e., the relative tetraploid content remained unchanged), depletion of TPX2, SGO1, CKAP5, AURKA, NIN, or KIF11 selectively suppressed tetraploids over diploids (i.e., WGD-selective; Fig. 1B). Among them, depletion of TPX2, a multifunctional mitotic regulator mediating microtubule nucleation, bundling, or stabilization (*32–35*), most strongly suppressed relative tetraploid content in the co-culture (Fig. 1B, S2A and B). Expression of an RNAi-resistant mCherry-TPX2 transgene restored the relative proliferation of tetraploids in TPX2-depleted co-culture, excluding possible off-target effects of RNAi (Fig. 1C and S2C).

We examined the effect of TPX2 depletion on mitotic progression in co-cultured GFP-labeled diploids and non-labeled tetraploids using live imaging (Fig. 1D and S2D; chromosomes were stained with SiR-DNA). In a mock-depleted condition, most diploids or tetraploids completed mitosis without defects. TPX2 depletion caused mitotic delay in both diploids and tetraploids. However, a significantly larger proportion of tetraploids showed extensive delay, with a higher frequency of mitotic death or chromosome segregation abnormalities than diploids (Fig. 1E, F, S2E, and F). Expression of RNAi-resistant mCherry-TPX2 rescued these mitotic defects in tetraploids, confirming RNAi specificity (Fig. 1G, H, S2G, and H). These results indicate that WGD confers increased TPX2 dependence for maintaining mitotic fidelity.

### WGD necessitates larger spindles to maintain their pole integrity

To examine potential WGD-linked effects of TPX2 depletion on mitotic spindle organization, we live-imaged the mitotic process in diploids and tetraploids expressing mCherry-α-tubulin, CENP-A-EGFP, and EGFP-GCP3 (labeling microtubules, centromeres, and centrosomes, respectively; Fig. 2A and S3A). Most diploids and tetraploids formed bipolar spindles and underwent equal chromosome segregation under mock depletion (Fig. 2A and B; Movie S1-4). TPX2 depletion differentially affected spindle organization between diploids and tetraploids: TPX2-depleted diploids frequently showed prolonged prometaphase with monopolar spindles, often followed by subsequent spindle bipolarization (Fig. 2A and B; Movie S2). On the other hand, TPX2-depleted tetraploids frequently manifested multipolar spindle defects (42%; Fig. 2A, B, and S3B; Movie S4). In most multipolar cells (90%), spindles became multipolar after bipolarization, with preformed poles split after chromosome alignment in TPX2-depleted tetraploids (Fig. 2B and S3B). The bipolar-to-multipolar conversion was much less frequent (2%) in their diploid counterparts, demonstrating that spindle pole integrity was attenuated in a WGD-specific manner under TPX2 depletion. We observed similar extents of WGD-dependent exacerbation of multipolar spindle defects in immunostaining analyses in HAP1, HCT116, and hTERT-RPE1 cellular backgrounds (Fig. S3C-H; to exclude a possible effect caused by supernumerary centrioles, we excluded cells with supernumerary centrioles from the analyses in Fig. S3E and H).

**Figure 2:**
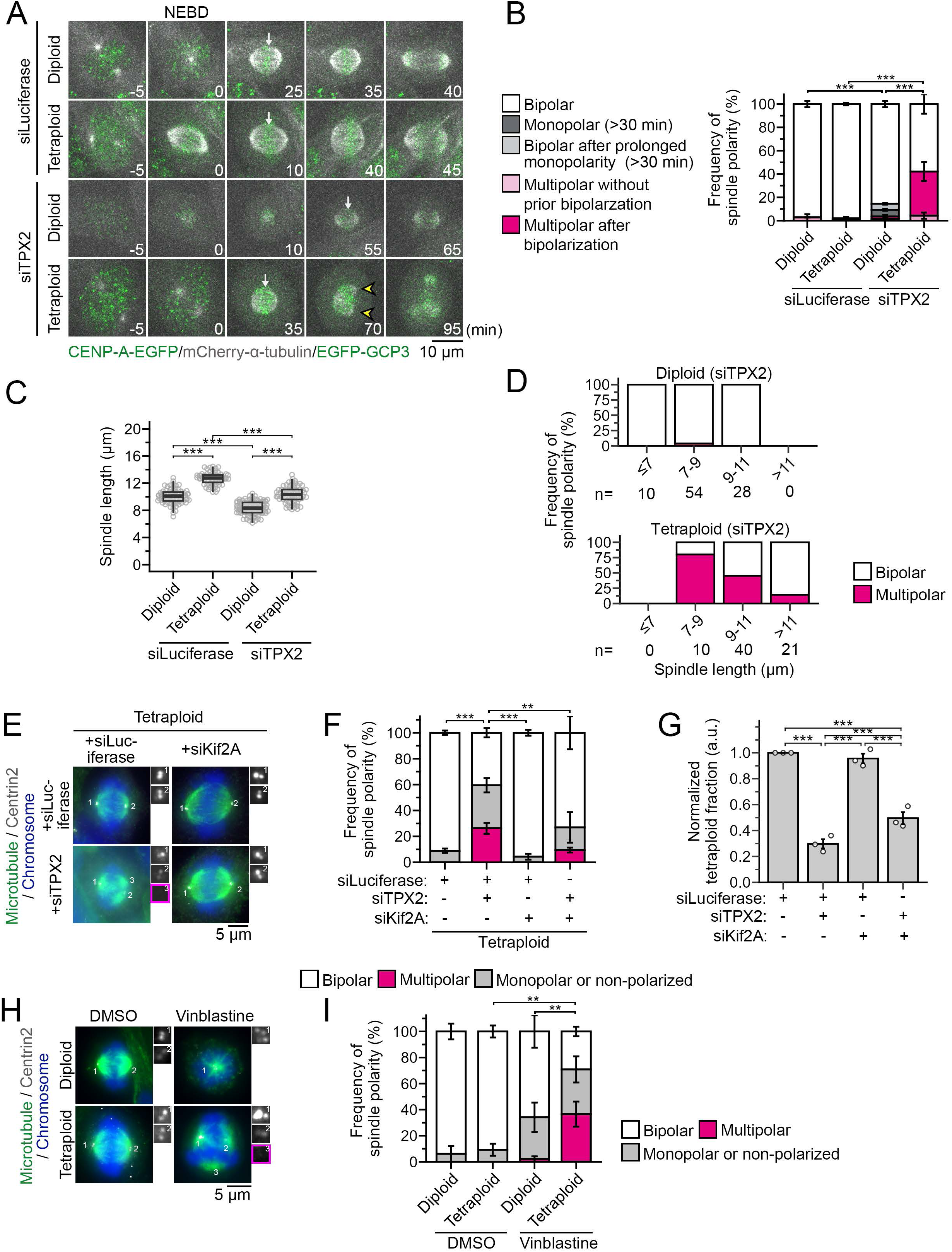
WGD necessitates larger spindles to maintain their pole integrity. **(A)** Time-lapse images of CENP-A-EGFP, mCherry-α-tubulin, and EGFP-GCP3 during mitosis in RNAi-treated diploids or tetraploids. Live images were initiated 18 h after siRNA transfection and taken at 5-min intervals. White arrows: cells that had just completed chromosome alignment. Yellow arrowheads: Splitting of the prefocused spindle poles. The timing of NEBD is set as 0 min. **(B)** Frequency of spindle polarity defects in A. Mean ± SE of 3 independent experiments. At least 83 cells from 3 independent experiments were analyzed for each condition. *** p<0.001; Fisher exact test. **(C)** Spindle length at the completion of chromosome alignment in A. Box plots show the median and the 25th and 75th percentiles. ***p<0.001; Tukey-Kramer test. **(D)** Frequency of bipolar-to-multipolar conversion, which was calculated for each bin of spindle length at the completion of chromosome alignment. At least 71 cells pooled from 3 independent experiments were analyzed for each condition in C and D. The number of cells analyzed for each length bin is shown at the bottom in D. **(E, H)** Immunostaining of α-tubulin and centrin2 in diploids or tetraploids treated with indicated siRNAs (F) or vinblastine (H). DNA was stained with DAPI. Insets: 3× images of centrin2 signal at respective spindle poles. Acentriolar poles are outlined in magenta. **(F, I)** Frequency of spindle polarity in E (F) or H (I). Mean ± SE of 3 independent experiments. At least 40 cells (F) or at least 24 cells (I), pooled from 3 independent experiments (only cells without supernumerary centrioles were analyzed). **p<0.01, ***p<0.001; Tukey-Kramer test following two-way ANOVA (applied only to multipolar spindle frequency). **(G)** Normalized tetraploid fraction in the RNAi-treated co-culture relative to the mock-depleted control. Mean ± SE of 3 independent experiments. ***p<0.001; Steel-Dwass test.

To address factors affecting spindle pole integrity under TPX2 depletion in tetraploids, we analyzed spindle sizes (length and width) in the live-imaging data above (Fig. 2C and S4A; to avoid mitotic stage-dependent bias, we measured sizes when chromosomes were fully aligned). In a mock-depleted condition, tetraploids form spindles significantly larger in length and width compared to diploids (13 µm vs 10 µm in length and 10 µm vs 7 µm in width, respectively; Fig. 2C and S4A), consistent with previous reports (*18–20, 36, 37*). TPX2 depletion significantly reduced spindle length and width in both diploids and tetraploids (Fig. 2C and S4A), as observed in other human cell lines (*36, 37*). In TPX2-depleted tetraploids, the frequency of multipolarization drastically increased, especially when spindle length was shorter than 9 μm (80% for spindle length ≤ 9 μm, whereas 14% for spindle length >11 μm; Fig. 2D). In contrast, most TPX2-depleted diploids kept spindle bipolarity even when they were ≤9 μm in length (Fig. S2D). The frequency of multipolarization was constant across different spindle width bins in TPX2-depleted tetraploids (Fig. S4B). This result indicates that WGD necessitates longer spindles to maintain bipolarity, suggesting an important role for WGD-linked spindle length scaling for mitotic fidelity.

We examined whether artificial spindle elongation was sufficient to cancel the increased TPX2 dependence in tetraploids. To this end, we co-depleted a spindle microtubule depolymerizer, Kif2A, whose depletion increased spindle length (*38–40*), in TPX-depleted tetraploids (Fig. 2E and S4C). Kif2A co-depletion significantly increased spindle length and restored spindle bipolarity in TPX2-depleted tetraploids (Fig. 2F and S4D; note that absolute values for spindle length and width were shorter in fixed samples compared to those in live-imaged samples due to sample shrinkage upon methanol fixation). Moreover, Kif2A co-depletion substantially improved the relative proliferation of tetraploids to diploids under TPX2 depletion (Fig. 2G and S4E). These data suggest that spindle shortening is the main cause of WGD-associated aggravation of mitotic and proliferative defects under TPX2 depletion. Consistent with this idea, orthogonal spindle downsizing by low-dose microtubule depolymerizers (nocodazole or vinblastine) also caused spindle multipolarization in a tetraploidy-selective manner (Fig. 2H, I, and S4F-H). Overall, these results show that WGD makes spindles more prone to multipolarization and confers dependence on larger spindles to maintain pole integrity.

### Multipolar spindle defect is the primary cause of the tetraploidy-selective suppression upon TPX2 loss

Different aspects of TPX2 function, including activation of Aurora A kinase, spindle microtubule bundling or nucleation, or regulation of Eg5 motor protein (*32, 33, 41, 42*), potentially contribute to spindle size and polarity control after WGD. Therefore, we examined the effects of TPX2 mutants in different domains (tagged with mCherry), expressed under endogenous TPX2 depletion, on spindle organization in tetraploids (Fig. S5A-C). Expression of mCherry-TPX2WT substantially resolved either monopolar or multipolar defects (Fig. S5C and D). Deletions or mutations in the nuclear localization signal sequences (5KMQ) (*43, 44*), the presumptive branching microtubule nucleation domains (4FA) (*32*), or the C-terminal end mediating its interaction with Eg5 (ΔC) (*42, 45, 46*) modestly reduced, albeit not completely abolished, the efficiency of the transgene in resolving the multipolar defects (Fig. S5D).

On the other hand, mCherry-TPX2ΔN, lacking the Aurora A-binding and activating domain (*37, 41*), solved multipolar defects as efficiently as mCherry-TPX2WT (Fig. S5A-D), while it failed to solve monopolar or non-polar defects (Fig. S5D). Consistently, non-phosphorylatable mutations on Aurora A-dependent sites on TPX2 (S121A/S125A) (*36*) or depletion of Aurora A had minimal effect on spindle multipolarization in tetraploids (Fig. S5A-G). We utilized the above property of mCherry-TPX2ΔN to assess cellular effects of suppressing multipolar defects without resolving other spindle defects. mCherry-TPX2ΔN expression substantially restored mitotic progression and relative proliferation of tetraploids (Fig. 1C, G, and H), demonstrating that the multipolar defects are the primary cause of tetraploidy-selective suppression upon TPX2 depletion. The data also indicate that Aurora A depletion selectively suppressed tetraploid cell proliferation through a different mechanism than TPX2 depletion (Fig. 1B), while the exact mechanism remains to be addressed in future studies.

### WGD increases mechanical stress on metaphase spindle poles via kinetochore-microtubule attachment

The tetraploidy-specific disintegration of the preformed spindle poles under TPX2 depletion (Fig. 2) raises the possibility that spindle poles endure stronger mechanical stress under the WGD condition. To directly address this idea, we conducted laser ablation microsurgery to cut a pole of a metaphase bipolar spindle and analyze the dynamics of spindle morphological changes in diploids and tetraploids expressing EGFP-α-tubulin and H2B-mCherry (Fig. 3A; cells were arrested at metaphase by MG132 treatment; Movie S5). Spindle pole ablation caused immediate recoiling of the ablated half spindle along its width, reflecting a mechanical stress imposed on the pole (Fig. 3A). The recoiling was significantly faster and more profound in tetraploids than in diploids (measured based on the distance between two cut edges of the ablated half spindle; Fig. 3B-D; Movie S5), indicating a stronger mechanical stress imposed on spindle poles after WGD.

**Figure 3:**
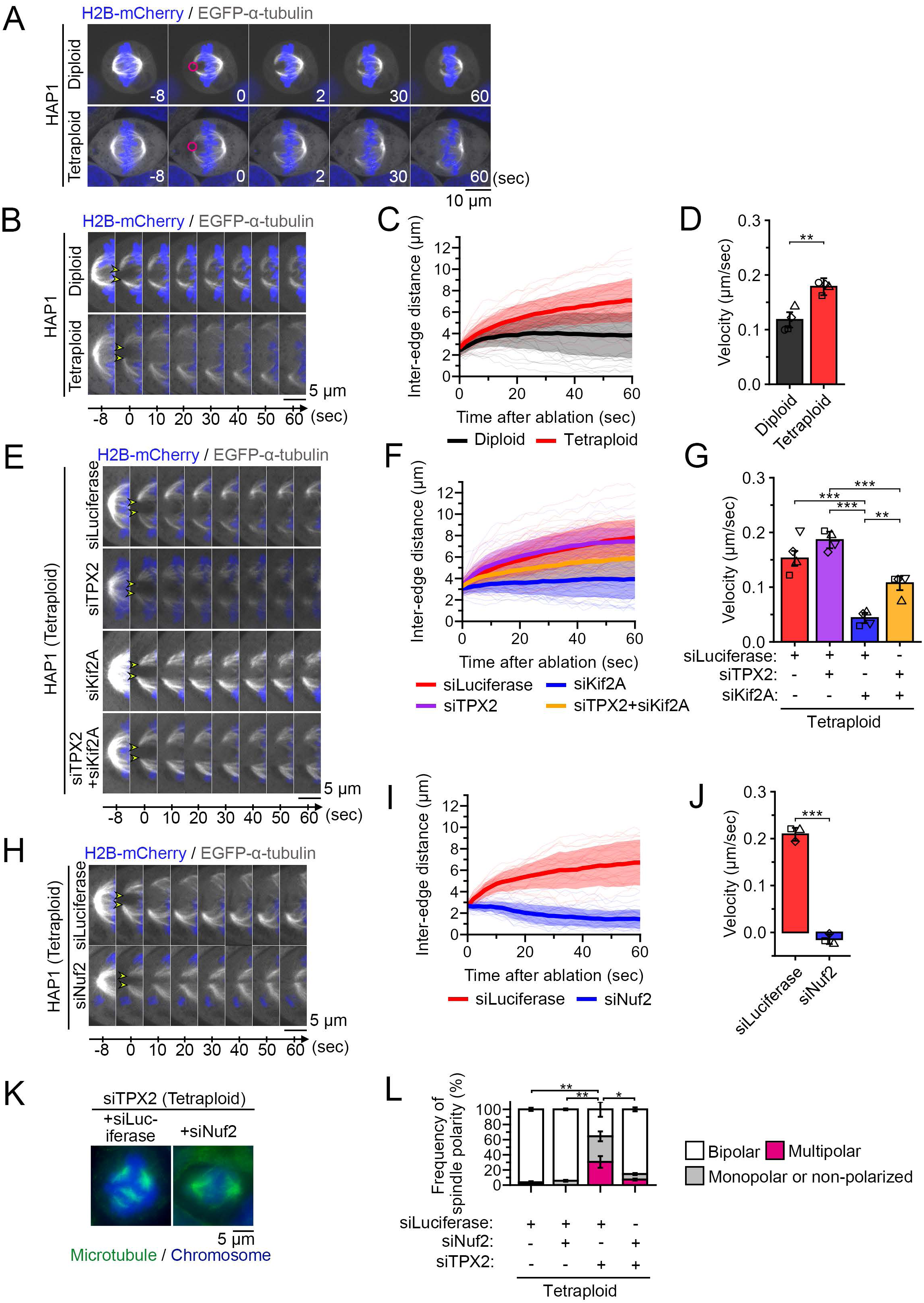
Increased mechanical stress at spindle poles in the tetraploid state. **(A, B, E, H)** Time-lapse images of EGFP-α-tubulin, and mCherry-H2B in diploids or tetraploids arrested at metaphase by MG132 treatment. Laser ablation was introduced at one of the spindle poles, as indicated by magenta circles in A. Cells were treated with the indicated siRNAs in E and H. Cropped views of the ablated side of the spindle are shown in B, E, and H. Cut edges of spindles are indicated by arrowheads. Live images were taken at 2-s intervals. The timing of laser ablation is set as 0 s. **(C, F, I)** Time course of the distance between two cut edges of the ablated side of the spindle in B (C), E (F), and H (I). Bold lines with shadows indicate the mean ± SD of at least 33 cells (C), at least 42 cells (F), or at least 27 cells (I) from at least 3 independent experiments. Fine lines indicate single-cell data. **(D, G, J)** Velocity of time-dependent changes in the inter-edge distance from 0 to 10 s after laser ablation in B (D), E (G), and H (J). Mean ± SE of at least 3 independent experiments. **p<0.01, ***p<0.001; unpaired t-test for D and J and Tukey-Kramer test for G. **(K)** Immunostaining of α-tubulin in RNAi-treated tetraploids. DNA was stained with DAPI. **(L)** Frequency of spindle polarity in K. Mean ± SE of 3 independent experiments. At least 56 cells were analyzed for each condition. *p<0.05, **p<0.01; Tukey-Kramer test following two-way ANOVA (applied only to multipolar spindle frequency).

To understand the impacts of spindle size on the mechanical stress at the spindle poles, we examined the dynamics of spindle recoiling upon laser ablation in tetraploids depleted of TPX2 and/or Kif2A (Fig. 3E-G, S6A, and Movie S6). While TPX2 depletion modestly facilitated recoiling with spindle shortening, Kif2A depletion significantly suppressed recoiling with spindle elongation (Fig. 3F and G). Among various characteristics of spindle structures (Fig. S6B-F), spindle length showed the strongest anticorrelation with recoiling rate across conditions (Fig. S6F and Supplementary Appendix 1). This result demonstrates that a larger spindle size alleviates the WGD-derived mechanical stress at the spindle poles, thereby securing robust spindle bipolarity upon drastic chromosome gain.

During the recoiling of laser-ablated spindles, the kinetochore fibers within the ablated half of the spindle moved outward in coordination with the paired kinetochore fibers on the opposite half of the spindle (Fig. S6G), consistent with previous observations of ablated kinetochore fibers (*47*). These data suggest mechanical coupling between paired kinetochore fibers. We therefore addressed whether kinetochore-microtubule attachment contributes to the transmission of the spindle pole stress in tetraploids. To this end, we analyzed spindle recoiling upon laser ablation in tetraploids depleted of Nuf2, which is essential for kinetochore-microtubule attachment (*48*). Nuf2 depletion almost completely suppressed the spindle recoiling upon ablation in tetraploids, demonstrating that the WGD-derived mechanical stress on the pole is applied in a manner depending on kinetochore-microtubule attachment (Fig. 3H-J; Movie S7 and S6A).

Based on the above results, we next examined the effects of co-depletion of Nuf2 on spindle organization in TPX2-depleted tetraploids (Fig. 3K, L, and S6H). Nuf2 co-depletion significantly suppressed multipolar spindle defects caused by TPX2 depletion and restored spindle bipolarity in tetraploids. Co-depletion of Mis12, another factor mediating kinetochore-microtubule attachment (*49*), also reverted spindle bipolarity in TPX2-depleted tetraploids (Fig. S6H-J). Therefore, WGD promotes multipolar spindle defects in a manner depending on kinetochore-microtubule attachment, indicating that spindle-chromosome interactions critically contribute to this structural vulnerability.

### Ploidy increase prevents bipolar spindle formation upon acute WGD

We next investigated how the above WGD-derived mechanical stress affects spindle fidelity upon de novo WGD, a process closely associated with cancer development, in which the doubled number of centrosomes interferes with the spindle polarity control (*5, 7*). We induced de novo WGD through cytokinesis failure in haploid, diploid, or tetraploid HAP1 cell lines (forming acute diploids, tetraploids, or octaploids, respectively) and examined spindle organization in the first mitosis after WGD (Fig. 4A and B). In acute diploids, 76% of cells formed bipolar spindles by clustering the excess centrosomes at the two poles (Fig. 4B and C), consistent with previous findings in other systems (*5, 6, 50*). Prevalence of bipolar spindles decreased as the ploidy level increased, rendering acute octaploids much less potent at forming bipolar spindles (12%; Fig. 4C). Nuf2 depletion increased the frequency of spindle bipolarization to 65% in acute octaploids (Fig. 4D, E, and S7A), demonstrating a profound contribution of kinetochore-microtubule attachment to mitotic instability upon de novo WGD in higher ploidy states.

**Figure 4:**
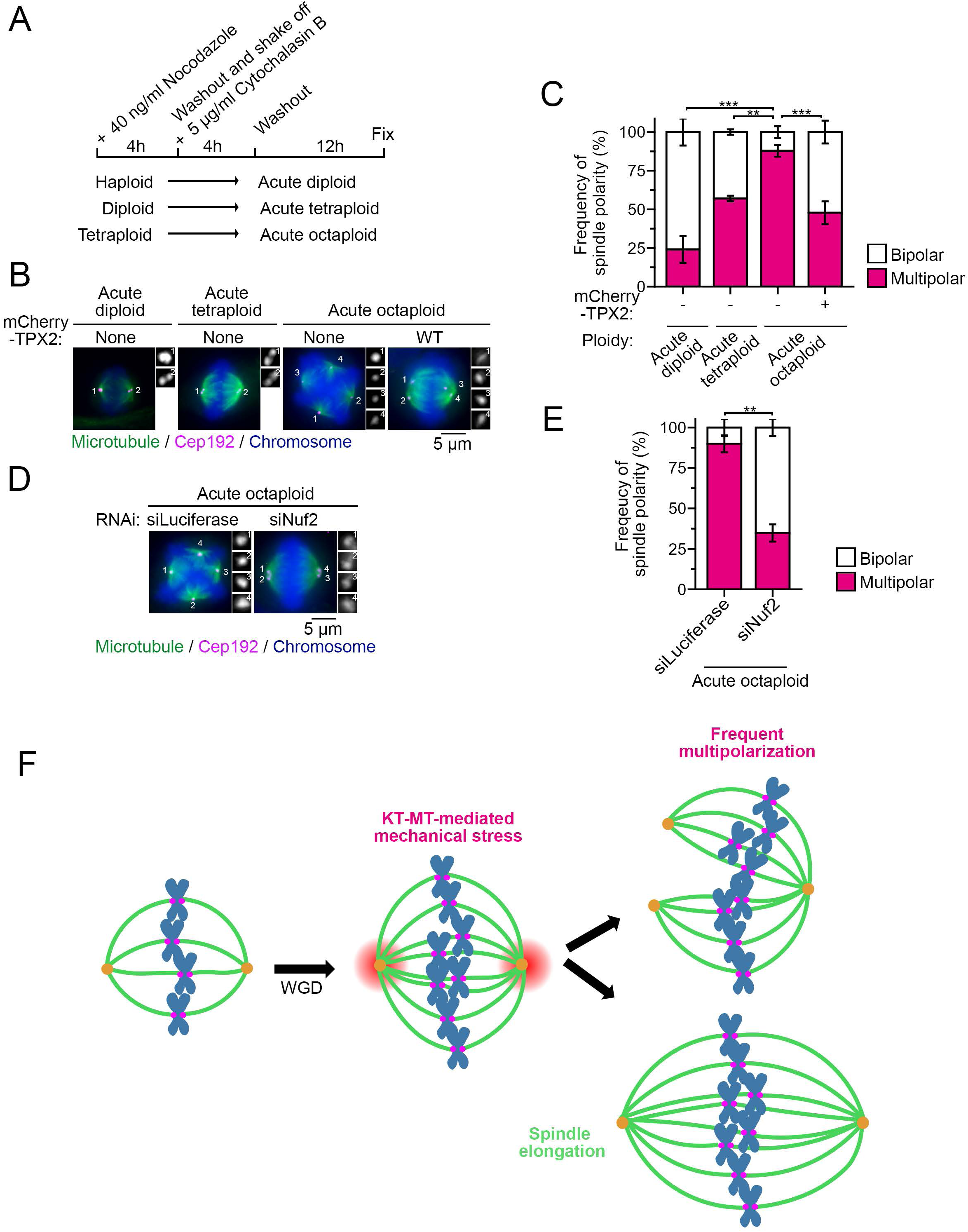
Chromosome increase interferes with spindle bipolarization upon acute WGD in a kinetochore-microtubule-dependent manner. **(A)** Scheme of acute WGD induction and centrosome clustering during the subsequent mitosis. **(B, D)** Immunostaining of α-tubulin and Cep192 in acute diploids, tetraploids, or octaploids. DNA was stained with DAPI. TPX2 WT-mCherry was expressed in the indicated sample in C. Cells were treated with the indicated siRNAs in D. **(C, E)** Frequency of spindle polarity in B (C) or D (E). Mean ± SE of 3 independent experiments. At least 35 cells (C) or at least 48 cells (E) in total for each condition. **p<0.01, ***p<0.001; Dunnett’s test (applied only to multipolar spindle frequency). **(F)** A model for increased polar mechanical stress and risk for multipolar defects associated with WGD.

Given TPX2’s essential contribution to bipolar spindle formation in chronic tetraploids (Fig. 2), we next examined whether TPX2 supplementation by transgenic expression of mCherry-TPX2 can improve spindle bipolarization in acute octaploids (Fig. 4B, C, and S7B). In chronic tetraploids, mCherry-TPX2 expression increased the net TPX2 protein level by 2.0-fold compared to the non-transfected control, resulting in increased accumulation of TPX2 on the spindle (by 1.2-fold) with a significant increase in spindle length (by 1.3-fold) (Fig. S7C-F). TPX2 supplementation substantially facilitated bipolar spindle formation in acute octaploids (Fig. 4B and C), demonstrating that TPX2-mediated size control, when expressed at a sufficiently high level, facilitates bipolar spindle formation even in acute WGD conditions at a ploidy state higher than tetraploidy.

## Discussion

In this study, we found that WGD increases mechanical stress at metaphase spindle poles, causing a higher risk of multipolar spindle defects in human cells. Multipolar spindle defects became particularly prevalent when spindle length fell below 9 μm, and artificial spindle elongation was sufficient to alleviate polar mechanical stress and multipolar defects in TPX2-depleted tetraploids (Fig. 2 and 3). These findings indicate that ploidy-linked spindle elongation plays an adaptive role in reducing the risk of multipolar defects and maintaining mitotic fidelity in post-WGD cells. Since the WGD-linked increase in polar mechanical stress depends on kinetochore-microtubule attachment, interactions between the doubled number of chromosomes and the spindle would contribute to the alterations in the spindle mechanical properties. Recent studies proposed that mitotic chromosomes that are packed into the metaphase plate push against each other, inducing the outward expansion of the plate upon WGD (*20, 51*). An intriguing possibility is that the WGD-associated outward expansion of the metaphase plate contributes to increasing polar mechanical stress via kinetochore fibers (Fig. 4F). Because the mechanics and force balances within the spindle are complex and not fully understood (*52, 53*), more systematic investigations in the future are required to elucidate the principles underlying the increased polar stress by WGD.

Previous studies have suggested that chromosome gain imposes physical constraints on mitosis after WGD, including interference with centrosome clustering and defects in chromosome alignment (*10, 11, 54*). Our findings extend this framework by identifying force transmission through kinetochore-microtubule attachment as a distinct physical challenge associated with increased chromosome content. Pleiotropic mitotic defects observed under WGD conditions should be interpreted in the context of these multiple chromosome-dependent physical constraints. In particular, we should consider that the multipolar defects upon acute WGD are partly caused by the centrosome-clustering barrier imposed by increased chromosome mass (*54*). However, the substantial rescue achieved by Nuf2 depletion indicates that chromosome-associated mechanical effects transmitted through kinetochore-microtubule attachment make an independent contribution to spindle instability at higher ploidy states. Thus, chromosome gain appears to compromise mitotic fidelity through multiple distinct physical mechanisms, among which force transmission through spindle microtubule architecture becomes increasingly important as chromosome content increases.

Our model predicts that kinetochore-microtubule-derived stress further increases with extremely high ploidy levels. Consistent with this view, the endogenous TPX2-dependent spindle-size control that supports bipolar spindle formation in tetraploids was insufficient in acute octaploids. This limited capacity to accommodate extreme chromosome gain may explain, at least in part, why proliferative cells with ploidy levels higher than octaploid are rare in animals. More broadly, our findings suggest that spindle architecture itself constrains the evolution of proliferative polyploidy: Highly focused spindle poles may render animal cells increasingly susceptible to chromosome-dependent mechanical stress as ploidy rises, whereas spindle architectures lacking discrete pole structures, like those in higher plants, could be less vulnerable to this limitation.

Our data also indicate that WGD-adaptive spindle size control is modulable, changing its functional level depending on TPX2 expression level (Fig. 4). One or more rounds of WGD are hallmarks of broad cancer types, potentially serving as a key pathway for malignant transformation (*55*). Interestingly, TPX2 expression shows a strong positive association with genome ploidy across cancer types (Fig. S8A and B) (*11*), indicating a possibility that cancer cells utilize the TPX2-dependent spindle size control to adapt to higher ploidy states after WGD. Therefore, the WGD-selective induction of mitotic failure observed in this study would provide an important clue for establishing WGD-targeting therapy across broad cancer types.

## Materials and methods

### Cell culture and flow cytometry

Cell lines and culture media used in this study are listed in Table S1. Media for all cell lines were supplemented with 10% fetal bovine serum (FBS) and 1× antibiotic-antimycotic solution (AA; Sigma-Aldrich). Cells were cultured at 37°C with 5% CO_2_. All experiments were performed with mycoplasma-free cells (tested using the MycoStrip mycoplasma detection kit; InvivoGen). We established transgenic cell lines by transfecting cells with plasmids encoding the corresponding transgenes using an Amaxa Nucleofector 2b device (Lonza), followed by selection of positive clones with the appropriate antibiotics listed in Table S1.

Flow cytometric analyses were conducted using a JSAN desktop cell sorter (Bay Bioscience). For live-cell DNA content analyses, cells were trypsinized using 0.05% trypsin-EDTA solution (Wako) and stained with 10 μg/ml Hoechst 33342 (Dojindo) for 15 min at 37°C. The stained cells were filtered through a cell strainer and applied to the cell sorter.

### Plasmids, compounds, and antibodies

The plasmid vectors constructed or used in this study, and the primers used for their construction are listed in Table S2 and Table S3, respectively. pmcherry-TPX2 or tdEos-CENPA-N-22 was a gift from Patricia Wadsworth (Addgene plasmid # 31227; RRID:Addgene_31227) (*45*) or Michael Davidson (Addgene plasmid # 57598; RRID:Addgene_57589). The siRNAs used in this study are listed in Table S4. Compounds and antibodies used in this study are listed in Table S5. siRNA transfection was performed using Lipofectamine RNAiMAX (Thermo Fisher Scientific).

### Live imaging

For live imaging, we seeded cells on an 8-well cover glass-bottom chamber (Iwaki or Zell-kontakt) coated with collagen Type I-C (Wako or Corning). We replaced the culture medium with phenol red-free Iscove’s Modified Dulbecco’s Medium (IMDM, Gibco) supplemented with 10% FBS and AA 1 h before imaging. For live-cell DNA staining, 200 nM SiR-DNA with 20 μM verapamil (Cytoskeleton, Inc.) was added to cell cultures 1 h before imaging. In all experiments, we analyzed only the first mitotic event from the start of live imaging. Cells were filmed at 37°C with 5% CO2 under a Ti2-E microscope (Nikon; equipped with a ×20 0.75 NA Plan-Apochromatic and Andor Zyla4.2 sCMOS camera), or a TE2000 microscope (Nikon; with a ×60 1.4 NA Plan-Apochromatic, Yokogawa CSU-X1 spinning-disc confocal unit, and Andor iXon3 EMCCD camera).

### Diploid-tetraploid co-culture experiment

We mixed and seeded EGFP-labeled diploid and non-labeled tetraploid cell suspensions in a 1:1 ratio (1.5 × 10^4^ cells/mL each, 2 mL for flow cytometry, and 1.25 × 10^4^ cells/ml each, 0.2 mL for live imaging). After 24 h, we transfected the co-cultures with siRNAs. For flow cytometry, we analyzed cells 48 h after transfection. We counted the two mixed cell populations separately based on EGFP fluorescence. Tetraploid cell proportion in each RNAi treatment was normalized to that in the mock-depleted (luciferase RNAi) control. For live imaging, we filmed cells from 48 h after transfection.

### Immunofluorescence staining

For immunostaining of centrin2, α-tubulin, and CEP192, cells were fixed with 100% methanol at −20°C for 10 min, and treated with BSA blocking buffer (150 mM NaCl, 10 mM Tris-HCl, pH 7.5, 5% BSA, and 0.1% Tween 20) at 25°C for 30 min. For immunostaining TPX2 and PCNT, cells were fixed with 3.2% paraformaldehyde in PBS at 25°C for 10 min, permeabilized with 0.5% Triton X-100 in PBS supplemented with 0.1 M glycine for 15 min at 25°C, and treated with 1% BSA in PBS. Fixed cells were then incubated with the primary antibodies overnight at 4°C, followed by incubation with the fluorescence-conjugated secondaries overnight at 4°C. DNA was stained with 1 μg/mL DAPI. Following each treatment, cells were washed 3 times with PBS.

Fixed cells were observed under a Ti2 microscope (Nikon; with a ×100 1.45 NA Plan-Apochromatic and Andor Zyla4.2 sCMOS camera), a Ti microscope (Nikon) with a ×100 1.40 NA Plan-Apochromatic, and Nikon A1Rsi (Nikon).

### Laser ablation experiment

For laser ablation, we used diploid or tetraploid HAP1 cells that expressed EGFP-α-tubulin (under a doxycycline promoter) and H2B-mCherry (under a constitutive EF1α promoter). Cells were seeded on a collagen-coated glass-bottom chamber (Iwaki) and treated with 1 µg/mL doxycycline 12.5 h before the initiation of imaging to induce EGFP-α-tubulin expression. The culture medium was then replaced with supplemented phenol red-free IMDM containing 3 μM MG132 30 min before imaging to avoid metaphase-anaphase transition during the experiment.

Laser ablation was performed using a 355 nm pulsed laser (2 kW, 1 uJ energy, 400 ps, 20 kHz) controlled by a galvanometric mirror on the iLAS3 system (Gataca Systems). Cells were imaged at 37°C with 5% CO2 under a Ti2-E microscope (Nikon) equipped with a x100 1.5 NA UPLAPO OHR oil-immersion objective, Yokogawa CSU-W1 spinning-disc confocal unit, and Oxxius LDI Prime laser-diode system comprising 405 nm (300 mW), 488 nm (1 W), 555 nm (1 W) and 640 nm (450 mW) lasers.

Dual-channel live imaging was performed using 488 nm and 561 nm lasers. Five pre-ablation images were acquired at 2-s intervals. Immediately following the pre-ablation sequence, 36-ms laser ablation pulse was applied using the 355-nm laser at 20% intensity to one of the bipolar spindle poles of metaphase cells using MetaMorph (Molecular Devices). Following ablation, 30 images were acquired at 2-s intervals. Local deformations of the ablated side of spindles were used as an indication of successful local ablation on the spindle structure.

### Quantitative analyses of imaging data

Quantification of spindle size (length and width) or distance between two cut edges of laser-ablated spindles was manually conducted using the segmented line tool in ImageJ (NIH). To quantify spindle morphology, we traced contours of the outermost kinetochore fibers using 26 roughly equidistant points with the multipoint tool in ImageJ. We coordinated the locations of the contour points so that points#1, 13, 14, and 26 corresponded to the chromosome-proximal edges of kinetochore fibers, while points#7 and 20 corresponded to the spindle poles (Fig. S6). We then fitted each spindle arc connecting the two spindle poles with a circle. The outermost spindle curvatures were then calculated as the reciprocal of the fitted circle radius, as previously described (*56*). The angles between two outermost kinetochore fibers flanking equatorial chromosomes (θ_equator_) or a spindle pole (θ_pole_) were calculated using a vector-based method. We represented the orientation of the two kinetochore fibers flanking the angle of interest as two vectors pointing toward the angle, based on 4 contour points along these fibers. Then, the angle between the 2 vectors was calculated from their normalized dot product according to

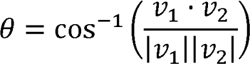

where *v*_1_ and *v*_2_ are vectors representing two kinetochore fibers, and *θ* represents the angle between these vectors.

In spindle polarity analyses, spindles with only one pole, or those with two poles with their distance less than 5 μm, were defined as “monopolar”. We manually quantified TPX2 immunostaining signal on spindles using the polygon selection tool in ImageJ. We performed background correction by subtracting the immunostaining signal within spindles from that in non-cell areas.

### Immunoblotting

Cells were lysed in SDS-PAGE sample buffer (1.025% SDS, 31.5 mM Tris-HCl, pH 6.8, 10.25% glycerol, 5% 2-mercaptoethanol) on ice, and then boiled for 5 min. Proteins separated by SDS-PAGE were transferred onto an Immun-Blot PVDF membrane (Bio-Rad). The membranes were blocked with 0.3% skim milk in Tween 20 Tris-buffered saline (TTBS; 50 mM Tris, 138 mM NaCl, 2.7 mM KCl, and 0.1% Tween 20), incubated with the primary antibodies overnight at 4°C, and incubated with horseradish peroxidase (HRP)-conjugated secondary antibodies for 1-3 h at 25°C. Each step was followed by 3 washes with TTBS. Signals were detected using the ezWestLumi plus ECL Substrate (ATTO) with a LuminoGraph II chemiluminescent imaging system (ATTO). Quantification of immunoblotting signals was conducted using the Gels tool in ImageJ. All uncropped images of immunoblots are shown in Supplementary Material S2.

### RT-qPCR

Approximately 0.5-1.0 × 10^6^ cells in asynchronous cultures were collected by trypsinization, and their total RNA was extracted using the NucleoSpin RNA kit (Macherey-Nagel). Reverse transcription was performed with 200 ng of total RNA in a 10 μL reaction using ReverTra Ace qPCR RT Master Mix (Toyobo). qPCR was then performed with 50 ng cDNA template and 500 nM forward and reverse primers using GeneAce SYBR^TM^ qPCR Mix II (Nippon Gene) on a Real-time PCR analysis system (Bio-Rad, CFX96). Each step was conducted according to the manufacturer’s instructions. We analyzed the data using the ΔΔCq method, with the mean Cq values of the β-actin gene used to calculate ΔCq. The primers used for qPCR are listed in Table S3.

### Bioinformatic analysis

The plot of gene expression levels in post-WGD tumors relative to non-WGD tumors was regenerated using tumor gene expression profile datasets and the analytical pipeline reported previously (*11*). The gene expression fold change and combined FDR values were plotted using the analysis procedures and parameters provided in the GitHub repository (https://github.com/campbio/Manuscripts/tree/master/Quinton_WGD_2020). The expression data for TPX2 were identified by querying the datasets, and the corresponding data point was highlighted within the regenerated plot.

For a comparative analysis of TPX2 expression levels between post-WGD and non-WGD cancer cell lines, TPX2 expression profiles were obtained from the DepMap public dataset (https://depmap.org/portal/api/download/files). Cell lines were classified as post-WGD (WGD^+^) or non-WGD (WGD^-^) based on the ABSOLUTE-derived WGD and ploidy calls reported previously (*11*).

### Statistical analysis

The normality of data distributions was assessed using the Shapiro-Wilk test, and homogeneity of variances was assessed using Levene’s test. Then, we chose an appropriate statistical test for each analysis based on data distribution and variance. Analyses for significant differences between the two groups were conducted using the unpaired Wilcoxon rank-sum test, unpaired Welch’s t-test, or unpaired or paired t-test. Multiple group analyses were conducted using the Tukey-Kramer test, the Fisher’s exact test with the Benjamini-Hochberg multiple testing correction, the Steel-Dwass test, or the Dwass-Steel Critchlow-Fligner (DSCF) test. Multiple group analyses with a common control were conducted using the Dunnett’s test or the Steel test. For spindle polarity analysis, the percentage of multipolar spindles was evaluated using Dunnett’s test or two-way ANOVA followed by the Tukey-Kramer test. All statistical analyses were conducted in R software. Statistical significance was set at p < 0.05. P-values are indicated in figures or the corresponding figure legends.

## Acknowledgment

We are grateful to Guang Yang, Ryoto Nomura, and Motonari Hamanaka for providing reagents; Iva Tolic, Nenad Pavin, Yusuke Toyama, and Renata Basto for useful discussions; the Global Facility Center at Hokkaido University for the flow cytometer and real-time PCR system; and the Nikon Imaging Center at Hokkaido University for microscopes.

## Author Contributions

Conceptualization, Q.W., T.Y., and R.U.; Methodology, Q.W., S.I-I., Y.N., T.Y., F.M., and R.U.; Investigation, Q.W., C.W., S.I-I., Y.N., and T.Y.; Formal Analysis, Q.W., C.W., T.Y., S.I-I., and R.U.; Resources, Q.W., Y.N., F.M., and R.U.; Supervision, R.U.; Writing – Original Draft, R.U.; Writing – Review & Editing, Q.W., Y.N., F.M., and R.U.; Funding Acquisition, Q.W., S.I-I., Y.N., F.M, and R.U.

## Funding and additional information

Q.W. and C.W. were supported by JST SPRING (JPMJSP2119). This work was supported by JSPS KAKENHI (Grant Numbers JP22KK0110, and JP23K19360 to S.I-I., JP21K19244, JP22H04926, JP24K21956, JP24KK0139, and JP24K02017 to R.U.), the Akiyama Life Science Foundation to S.I-I., the Princess Takamatsu Cancer Research Fund, the Orange Foundation, the Smoking Research Foundation, Daiichi Sankyo Foundation of Life Science, the Akiyama Life Science Foundation, the Hoansha Foundation, Sumitomo Electric Group CSR Foundation, and the Terumo Life Science Foundation to R.U.

**Supplementary Materials S1:** Results of the co-culture RNAi screen Flow cytometric analysis of EGFP and Hoechst intensity in co-cultures under all RNAi conditions, associated with Fig. 1A-B. Tetraploid proportion in each co-culture is indicated within the graph.

**Supplementary Materials S2:** Original immunoblot membranes used in this study Uncropped images of all immunoblots are shown, with the approximate indications of the cropped regions.

**Movie S1:** Mitosis in a mock-depleted diploid cell Live imaging of CENP-A-EGFP, mCherry-α-tubulin, and EGFP-GCP3 during mitosis in a mock-depleted diploid cell. Images were taken at 5-min intervals. The timing of NEBD is set as 0 min. The movie is shown at 900x real-time.

**Movie S2:** Mitosis in a TPX2-depleted diploid cell Live imaging of CENP-A-EGFP, mCherry-α-tubulin, and EGFP-GCP3 during mitosis in a TPX2-depleted diploid cell. Images were taken at 5-min intervals. The timing of NEBD is set as 0 min. The movie is shown at 900x real-time.

**Movie S3:** Mitosis in a mock-depleted tetraploid cell Live imaging of CENP-A-EGFP, mCherry-α-tubulin, and EGFP-GCP3 during mitosis in a mock-depleted tetraploid cell. Images were taken at 5-min intervals. The timing of NEBD is set as 0 min. The movie is shown at 900x real-time.

**Movie S4:** Mitosis in a TPX2-depleted tetraploid cell Live imaging of CENP-A-EGFP, mCherry-α-tubulin, and EGFP-GCP3 during mitosis in a TPX2-depleted tetraploid cell. Images were taken at 5-min intervals. The timing of NEBD is set as 0 min. The movie is shown at 900x real-time.

**Movie S5:** Laser ablation in mitotic diploids and tetraploids Live imaging of EGFP-α-tubulin and mCherry-H2B in diploids or tetraploids arrested at metaphase by MG132 treatment. Laser ablation was introduced at one of the spindle poles (the left pole) at time point 0 s. Live images were taken at 2-s intervals. The movie is shown at 20x real-time.

**Movie S6:** Laser ablation in RNAi-treated mitotic tetraploids Live imaging of EGFP-α-tubulin and mCherry-H2B in mock-, TPX2-, Kif2A-, or TPX2- and Kif2A-depleted diploids or tetraploids arrested at metaphase by MG132 treatment. Laser ablation was introduced at one of the spindle poles (the left pole) at time point 0 s. Live images were taken at 2-s intervals. The movie is shown at 20x real-time.

**Movie S7:** Laser ablation in Nuf2-depleted mitotic tetraploids Live imaging of EGFP-α-tubulin and mCherry-H2B in mock-or Nuf2-depleted diploids or tetraploids arrested at metaphase by MG132 treatment. Laser ablation was introduced at one of the spindle poles (the left pole) at time point 0 s. Live images were taken at 2-s intervals. The movie is shown at 20x real-time.

---

**Figure S1:**
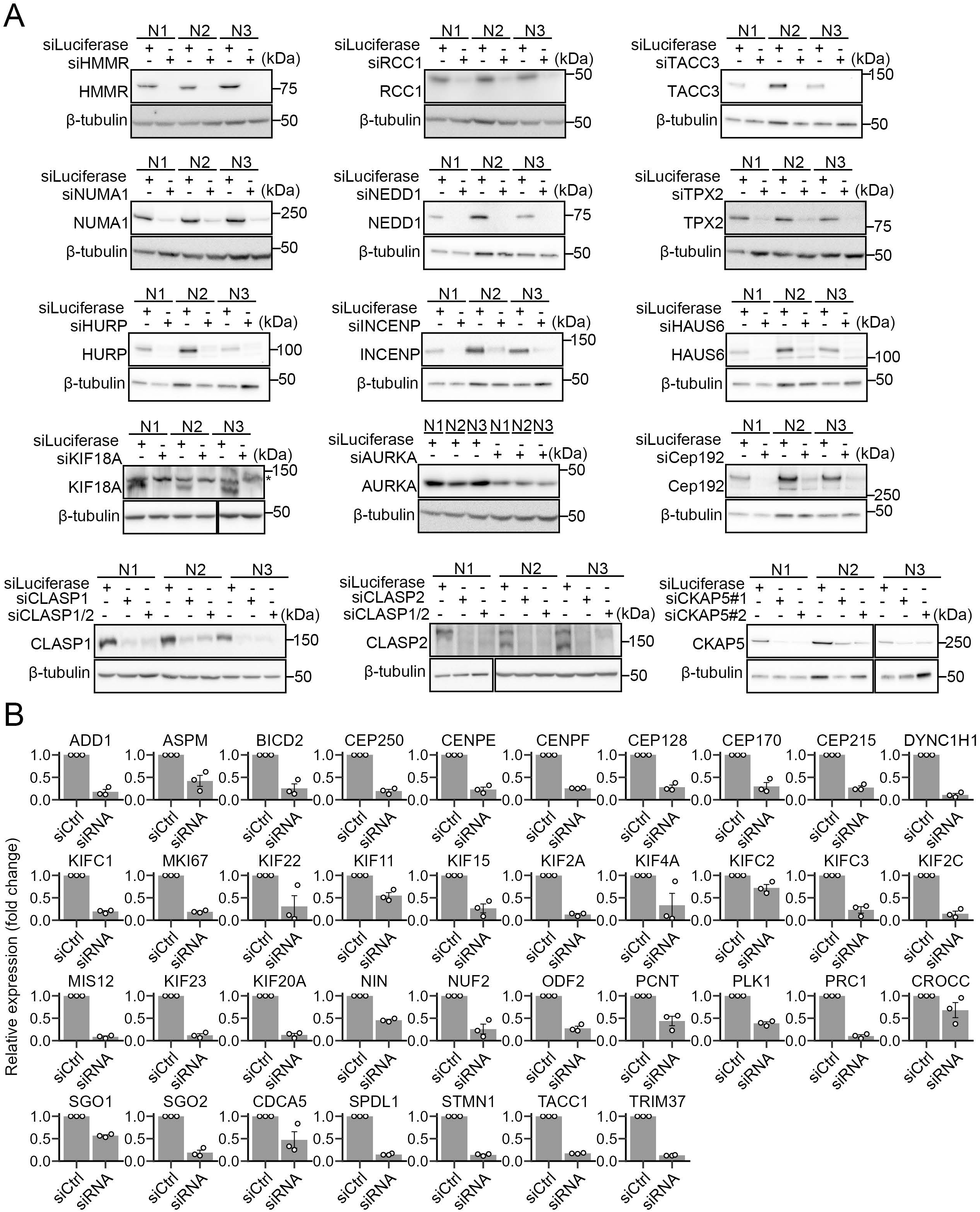
Confirmation of silencing of target genes in RNAi screen. **(A)** Immunoblotting of RNAi target proteins in diploid-tetraploid co-cultures treated with different siRNAs. β-tubulin was detected as a loading control. **(B)** Relative expression levels of RNAi target genes (normalized to that of β-actin) quantified by qPCR. Mean ± SE of 3 independent experiments.

**Figure S2:**
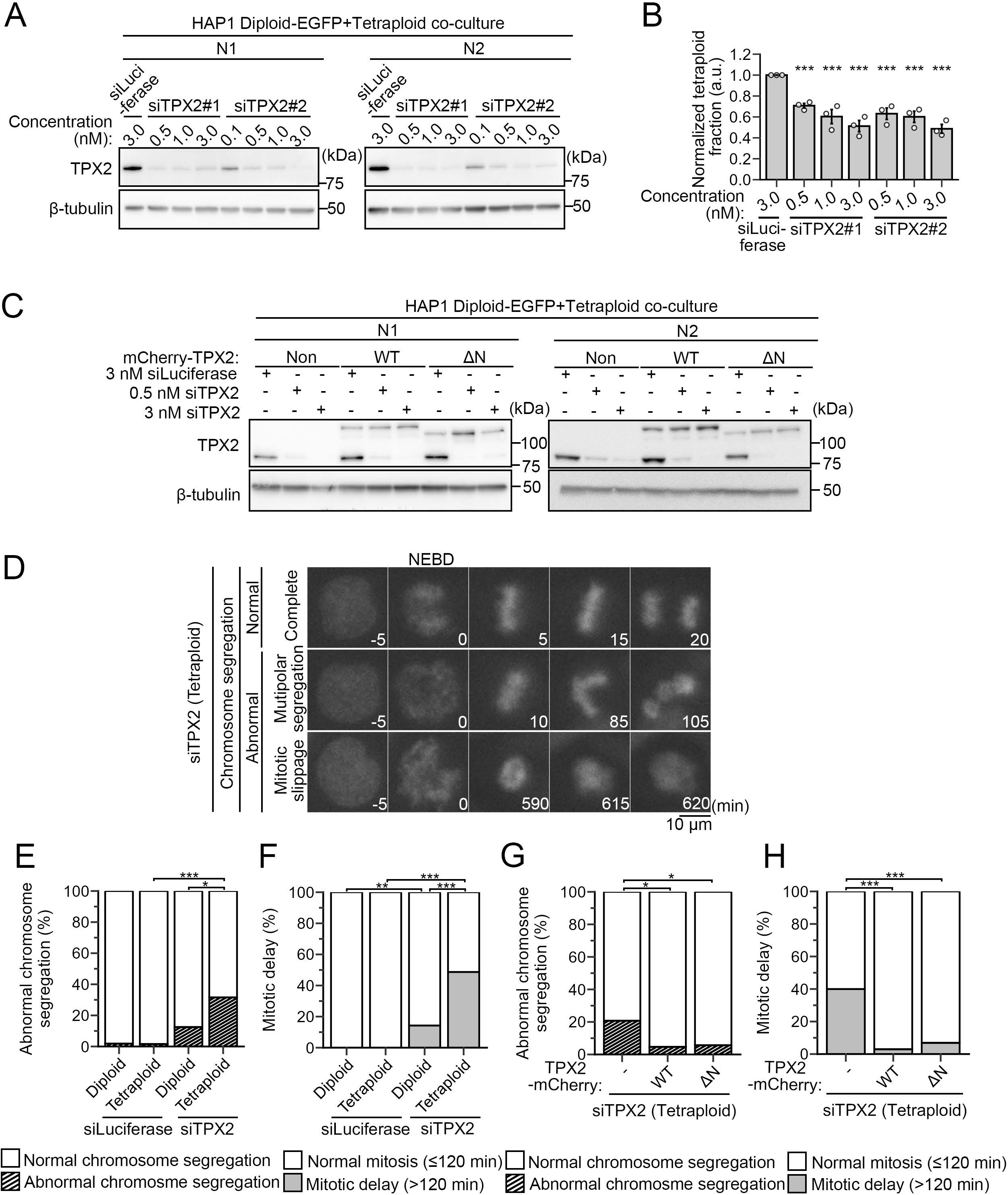
Profiling of TPX2-depleted tetraploid HAP1 cells. **(A, C)** Immunoblotting of TPX2 in diploid-tetraploid co-cultures with the indicated backgrounds, treated with different siRNAs targeting TPX2. β-tubulin was detected as a loading control. **(B)** Normalized tetraploid fraction in diploid-tetraploid co-culture treated with different siRNAs targeting TPX2 (relative to the mock-depleted control). Mean ± SE of 3 independent experiments. ***p<0.001; Steel test. **(D)** Time-lapse images of Sir-DNA-stained chromosomes during mitosis in TPX2-depleted tetraploids. Different examples of live imaging analysis are shown in Fig. 1D. The timing of NEBD is set as 0 min. **(E-H)** Frequency of abnormal chromosome segregation (E, G) or frequency of drastic mitotic delay (> 120 min; within the cells that completed mitosis; F, H). These analyses correspond to data in Fig. 1D (E, F) or Fig. 1G (G, H). At least 53 cells (E), 58 cells (G), 53 cells (F), or 66 cells (H), pooled from 2 independent experiments, were analyzed for each condition. *p<0.05, **p<0.01, ***p<0.001; Fisher exact test.

**Figure S3:**
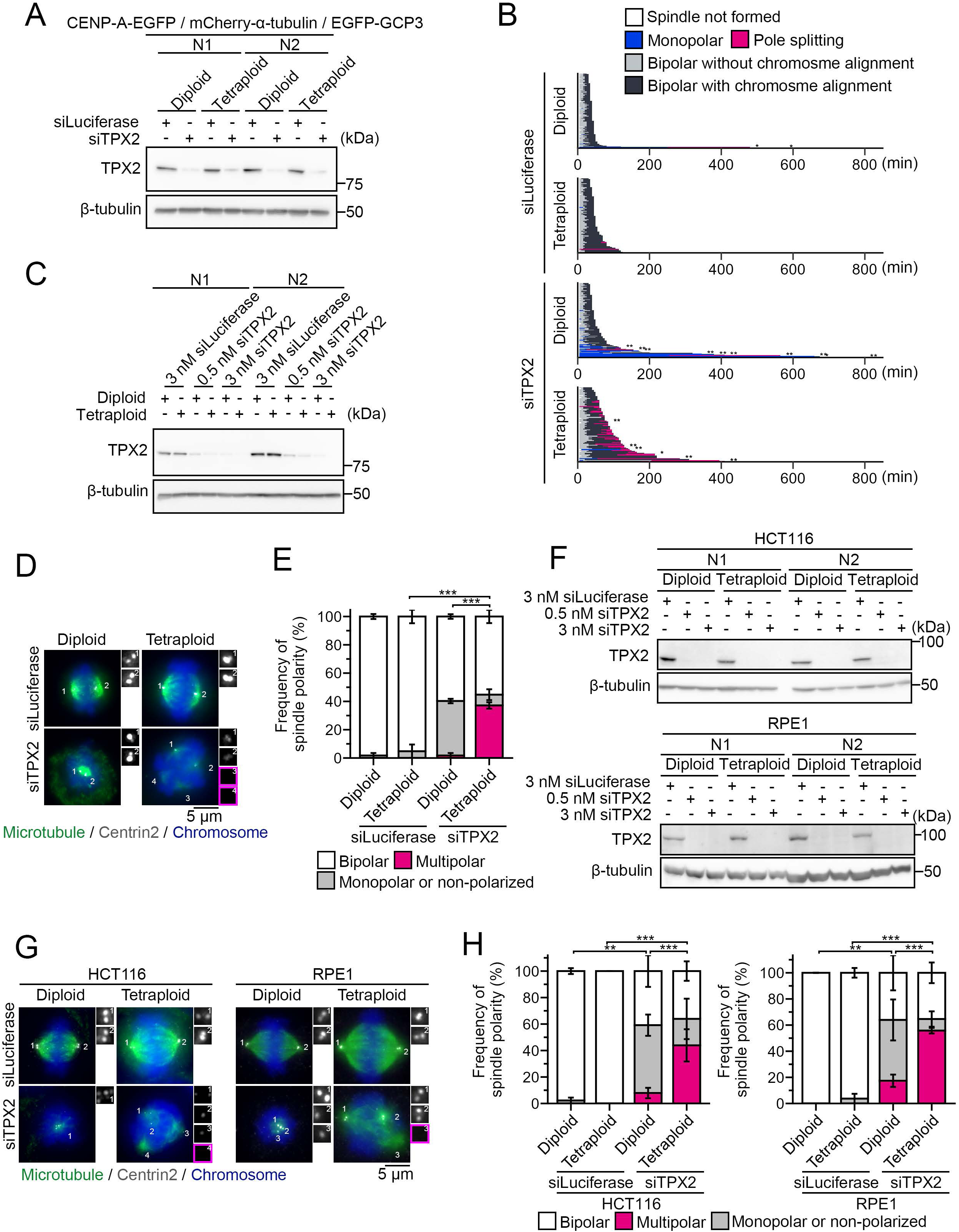
Tetraploidy-linked aggravation of multipolar defects upon TPX2 depletion across different cellular backgrounds. **(A)** Immunoblotting of TPX2 in RNAi-treated diploids or tetraploids in live imaging in Fig. 2A. β-tubulin was detected as a loading control. **(B)** Mitotic progression in live imaging in Fig. 2A. Each bar represents a single mitotic event (from NEBD to mitotic exit or the end of imaging) in a dividing cell. At least 83 cells from at least 3 experiments were analyzed for each condition. Asterisks indicate cells that underwent mitotic death, and double asterisks indicate the cells in which mitotic events could not be traced until mitotic exit. **(C, F)** Immunoblotting of TPX2 in RNAi-treated diploids or tetraploids in different cellular backgrounds. β-tubulin was detected as a loading control. **(D, G)** Immunostaining of α-tubulin and centrin2 in RNAi-treated diploid or tetraploid HAP1 (D), HCT116, or RPE1 cells (G). DNA was stained with DAPI. Insets: 3× images of centrin2 signal at respective spindle poles. Acentriolar poles are outlined in magenta. **(E, H)** Frequency of spindle polarity in D (E) or G (H). Mean ± SE of 3 independent experiments. At least 49 cells (E) or at least 24 cells (H) were analyzed for each condition. Only cells without supernumerary centrioles were analyzed. **p<0.01, ***p<0.001; Tukey-Kramer test following two-way ANOVA (applied only to multipolar spindle frequency).

**Figure S4:**
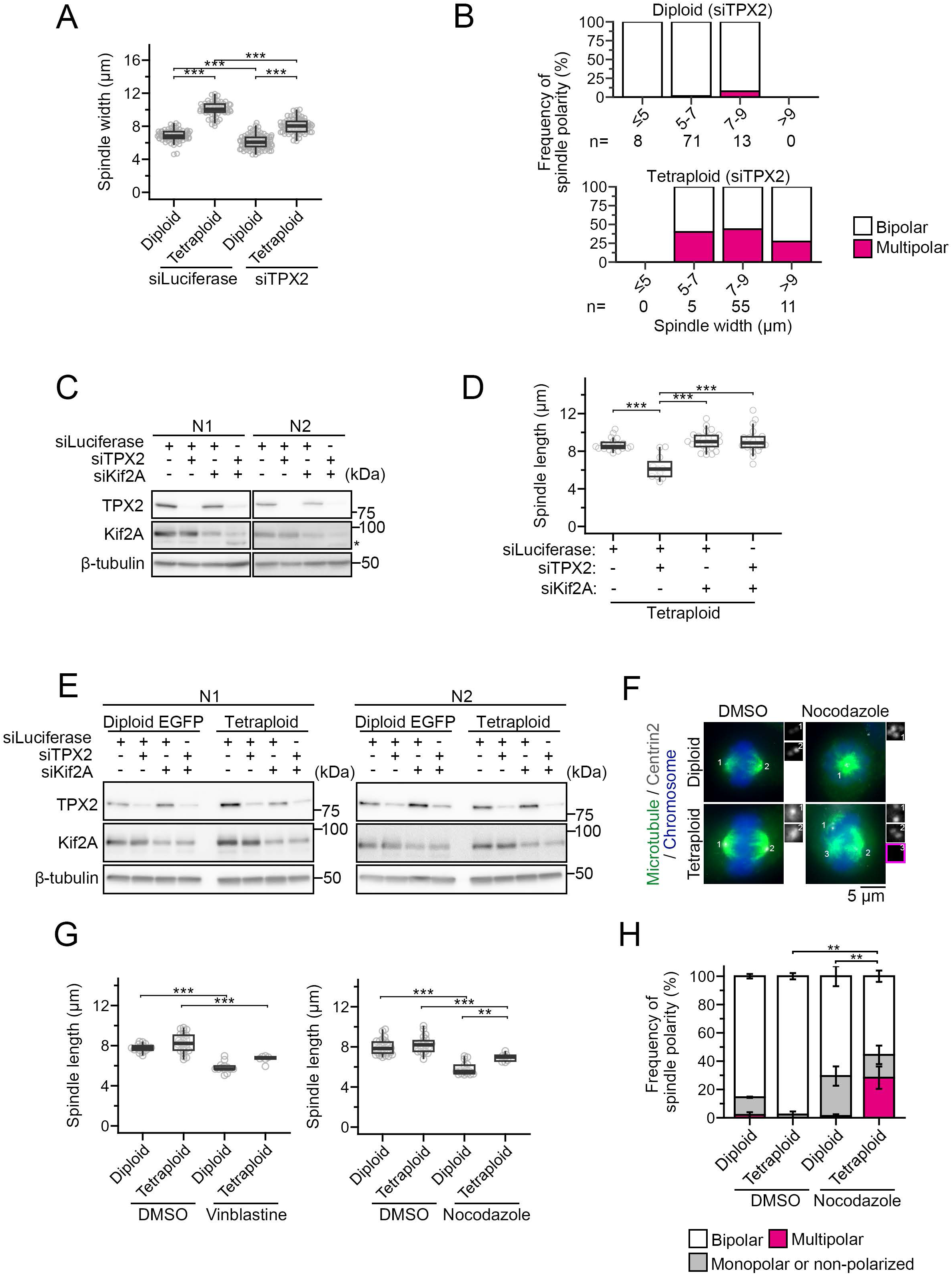
Spindle size quantification in different experimental conditions. **(A)** Spindle width at the completion of chromosome alignment in Fig. 2A. Box plots indicate the median, the 25th and 75th percentiles. ***p<0.001; Tukey-Kramer test. **(B)** Frequency of bipolar-to-multipolar conversion, which was calculated for each bin of spindle width at the completion of chromosome alignment. At least 71 cells pooled from at least 3 independent experiments were analyzed for each condition in A and B. The number of cells analyzed for each length bin is shown at the bottom in B. **(C, E)** Immunoblotting of TPX2 or Kif2A in RNA-treated tetraploid HAP1 cells used for the immunofluorescence (C) and co-culture flow cytometry (E) experiments, respectively. β-tubulin was detected as a loading control. **(D, G)** Quantification of spindle length (pole-to-pole in bipolar spindles) in RNAi-treated tetraploids (D) or compound-treated diploids or tetraploids (G). Box plots indicate the median, the 25^th^ and 75^th^ percentiles. At least 11 cells (D) or at least 6 cells (G), pooled from 3 independent experiments. **p<0.01, ***p<0.001; Tukey-Kramer test. **(F)** Immunostaining of α-tubulin and centrin2 in diploids or tetraploids treated with nocodazole. DNA was stained with DAPI. Insets: 3× images of centrin2 signal at respective spindle poles. Acentriolar poles are outlined in magenta. **(H)** Frequency of spindle polarity in F. Mean ± SE of 3 independent experiments. At least 48 cells were analyzed for each condition. Only cells without supernumerary centrioles were analyzed. **p<0.01; Tukey-Kramer test following two-way ANOVA (applied only to multipolar spindle frequency).

**Figure S5:**
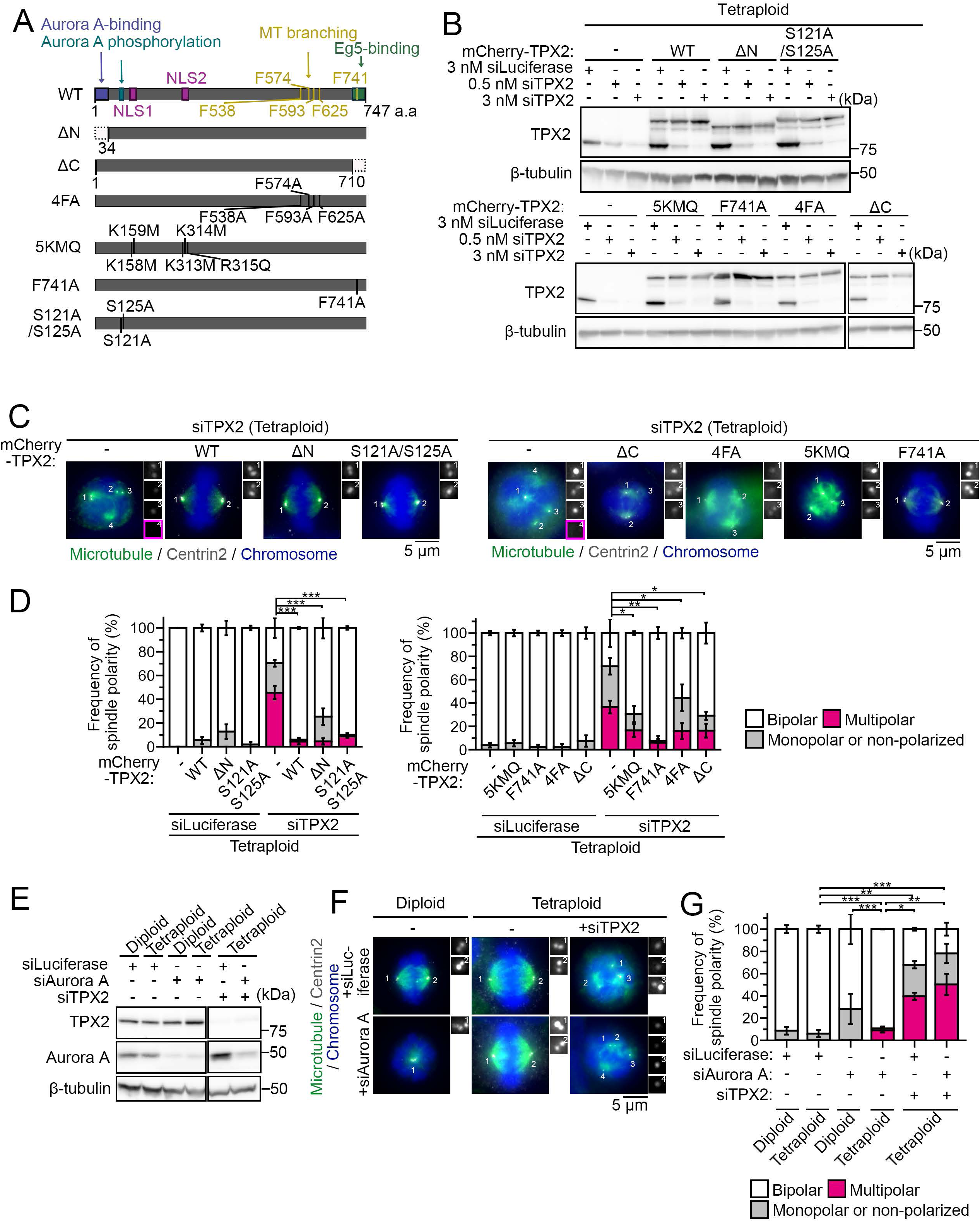
Effects of different TPX2 mutations or Aurora A depletion on spindle organization in tetraploids. **(A)** Scheme of TPX2 mutants used in the experiment. **(B, E)** Immunoblotting of TPX2 in RNAi-treated tetraploids expressing different TPX2 mutants (B) or Aurora A in RNAi-treated diploids or tetraploids (E). β-tubulin was detected as a loading control. **(C, F)** Immunostaining of α-tubulin and centrin2 in RNAi-treated tetraploids expressing different TPX2 mutants (C) or RNAi-treated diploids or tetraploids (F). DNA was stained with DAPI. Insets: 3× images of centrin2 signal at respective spindle poles. Acentriolar poles are outlined in magenta. **(D, G)** Frequency of spindle polarity in C (D) or F (G). Mean ± SE of 3 independent experiments. At least 40 cells (D) or at least 43 cells (G) were analyzed in total for each condition. Only cells without supernumerary centrioles were analyzed. *p<0.05, **p<0.01, ***p<0.001; Tukey-Kramer test following two-way ANOVA (applied only to multipolar spindle frequency).

**Figure S6:**
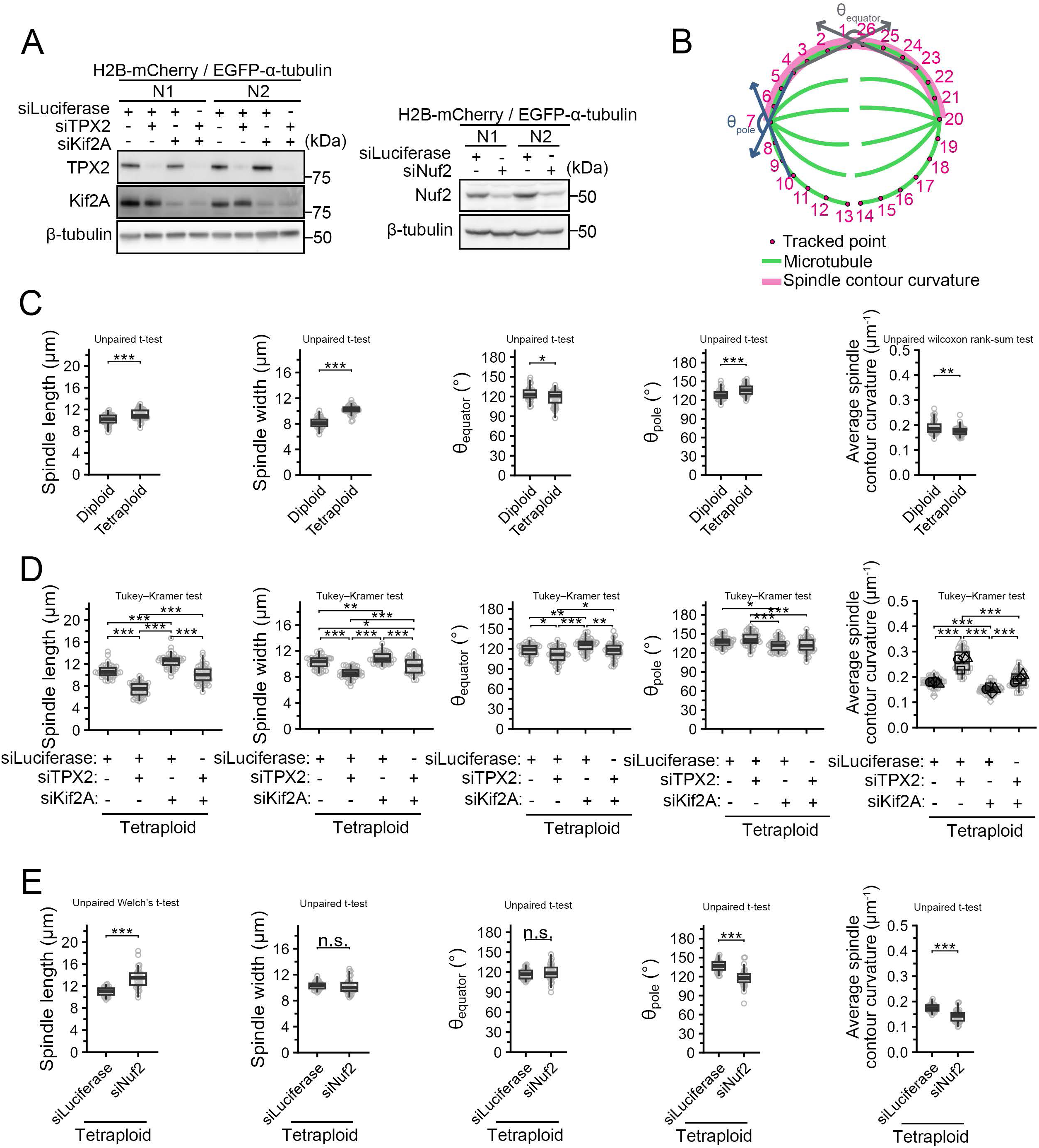
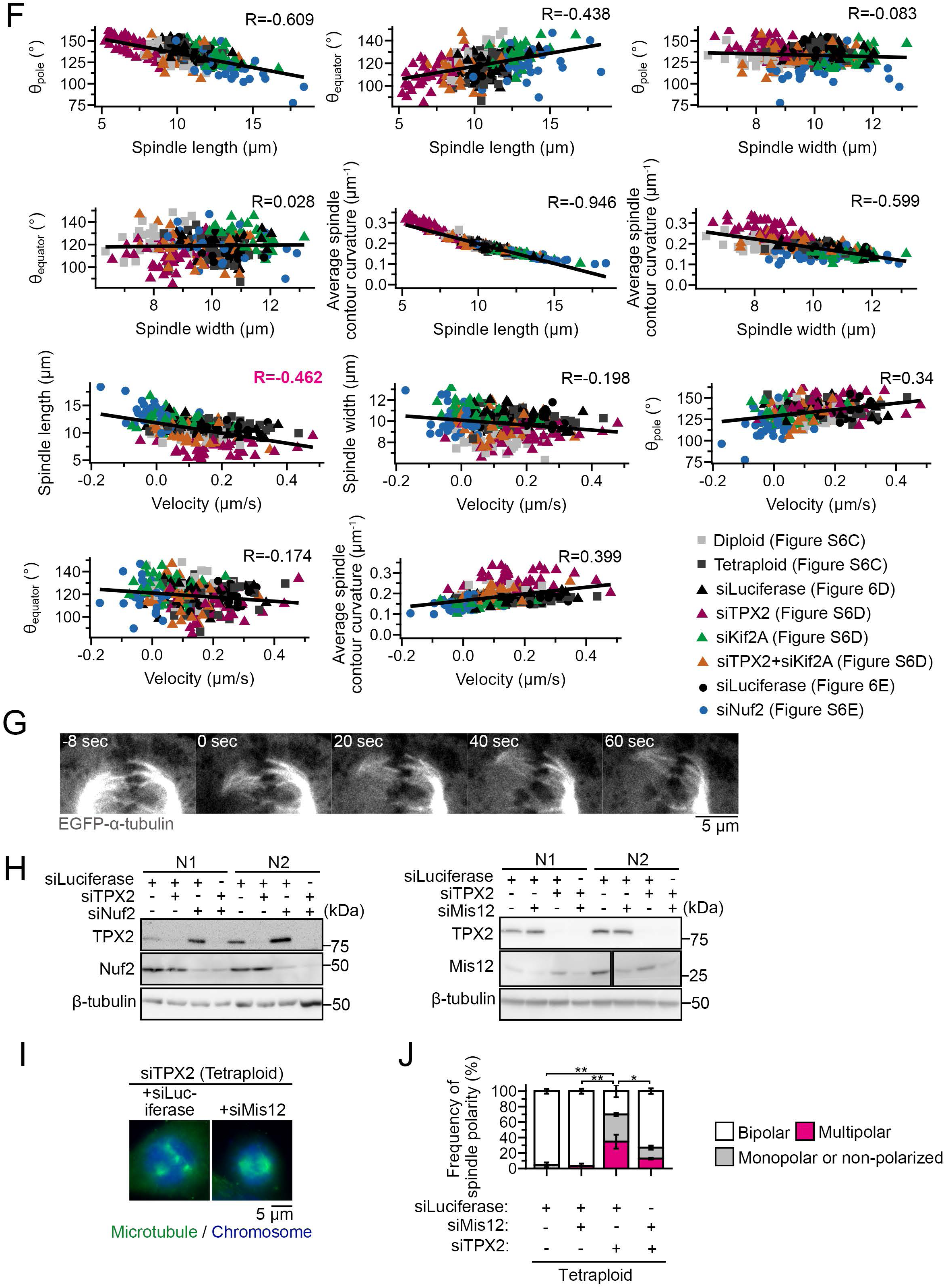
Confirmation of protein depletion in co-depletion experiment. **(A)** Immunoblotting of Nuf2 or Kif2A in RNA-treated tetraploid HAP1 cells used for the laser ablation experiments, respectively. β-tubulin was detected as a loading control. **(B)** Scheme of quantification of spindle morphological features. The indicated values were quantified in the pre-ablated spindles in Fig. 4B, E, and H. **(C-E)** Comparison of spindle size and morphological features between diploids and tetraploids (C) or among different RNAi conditions in tetraploids (D and E). Box plots indicate the median and the 25th and 75th percentiles. At least 33 cells (C), at least 42 cells (D), or at least 27 cells (E), pooled from at least 3 independent experiments. *p<0.05, **p<0.01, ***p<0.001; applied statistical analyses are indicated in the graphs. **(F)** Two-dimensional plots between indicated quantitative values. Data points from C-E are integrated into one plot. The Pearson correlation coefficient (R) was calculated across all experimental conditions. Linear regression lines were fitted to the pooled data and shown without confidence intervals. **(G)** A demonstration of co-migration of paired kinetochore fibers in opposite half spindles after laser ablation. **(H)** Immunoblotting of RNAi target proteins in tetraploids treated with different siRNAs. β-tubulin was detected as a loading control. **(I)** Immunostaining of α-tubulin in RNAi-treated tetraploids. DNA was stained with DAPI. **(J)** Frequency of spindle polarity in I. Mean ± SE of 3 independent experiments. At least 56 cells were analyzed for each condition. *p<0.05, **p<0.01; Tukey-Kramer test following two-way ANOVA (applied only to multipolar spindle frequency).

**Figure S7:**
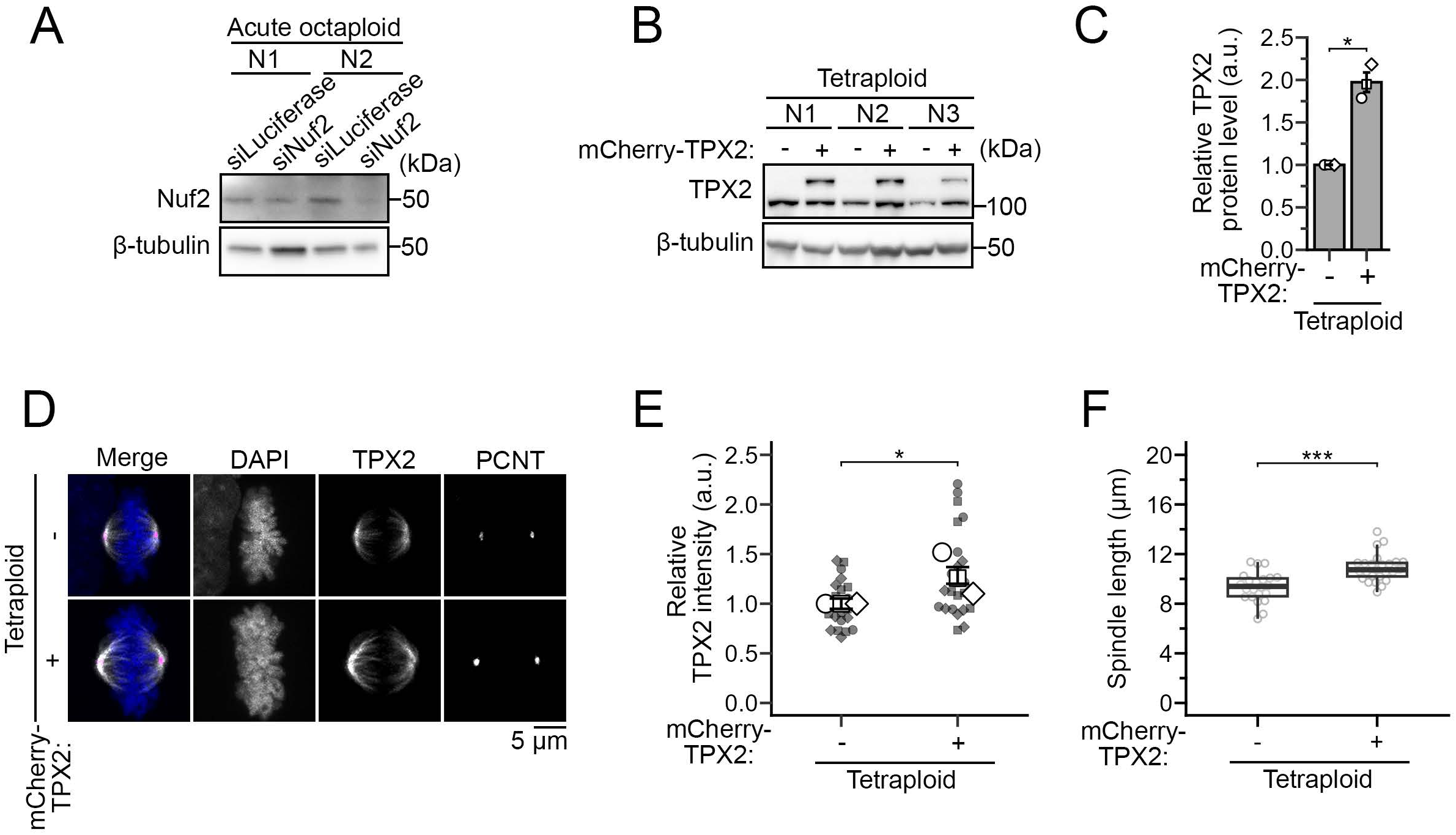
Supplementary data for acute octaploidization experiments. **(A, B)** Immunoblotting of Nuf2 in RNAi-treated acute octaploids (A) or TPX2 in chronic tetraploids with different backgrounds (B). β-tubulin was detected as a loading control. **(C)** Quantification of TPX2 expression levels in B. Mean ± SE of 3 independent experiments. *p<0.05; paired t-test. **(D)** Immunostaining of TPX2 and PCNT in tetraploids with or without exogenous expression of TPX2-mCherry. DNA was stained with DAPI. **(E, F)** Quantification of TPX2 immunostaining intensity on the spindle (E) or spindle length (pole-to-pole in bipolar spindles; F). Mean ± SE of 3 independent experiments in E (at least 22 cells were analyzed in total for each condition). Box plots indicate the median and the 25th and 75th percentiles in F (at least 22 cells pooled from 3 independent experiments). *p<0.05, ***p<0.001; unpaired Wilcoxon rank-sum test for E and unpaired t-test for F.

**Figure S8:**
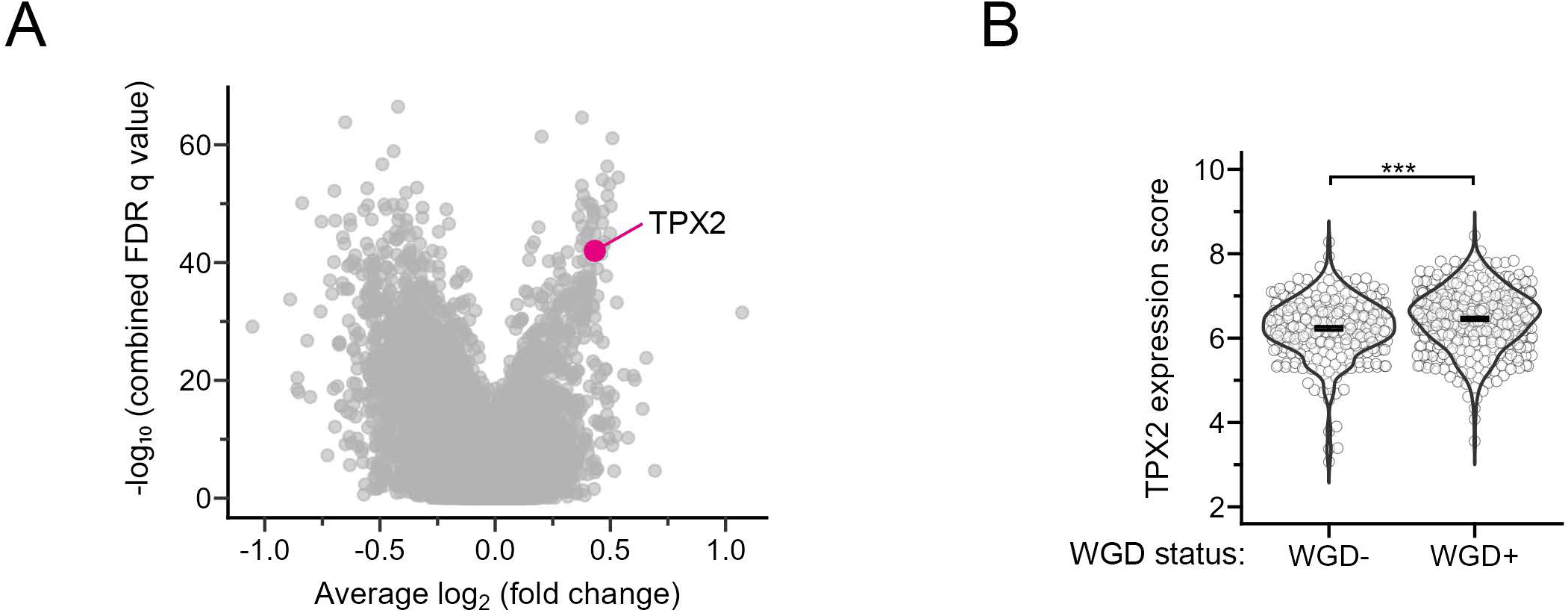
Increased TPX2 expression in post-WGD tumor cells. **(A)** A plot of relative expression levels of genes in WGD^+^ (post-WGD) tumors relative to WGD^-^ (non-WGD) tumors, which was regenerated using pancancer gene expression profiles and the analytical pipeline established and reported previously (*11*) (see Materials and methods for details; note that the distribution of data points is therefore identical to Fig. 2C in Quinton et al., 2021, but the TPX2 gene is highlighted here). Fold changes of gene expression in WGD^+^ tumors relative to WGD^-^ (non-WGD) tumors, plotted against combined FDR q-values across all tumor types. **(B)** TPX2 expression in WGD^-^ and WGD^+^ cell lines. TPX2 expression data were obtained from the DepMap OmicsExpressionProteinCodingGenesTPMLogp1_25Q2 dataset. WGD status was determined according to the WGD/Ploidy calls reported previously (*11*). ***p<0.001, unpaired Wilcoxon rank-sum test.

