## Supplementary figures and images for "Spindle size scaling secures mitotic fidelity against whole-genome doubling-associated mechanical stress"

### Supplementary material S1

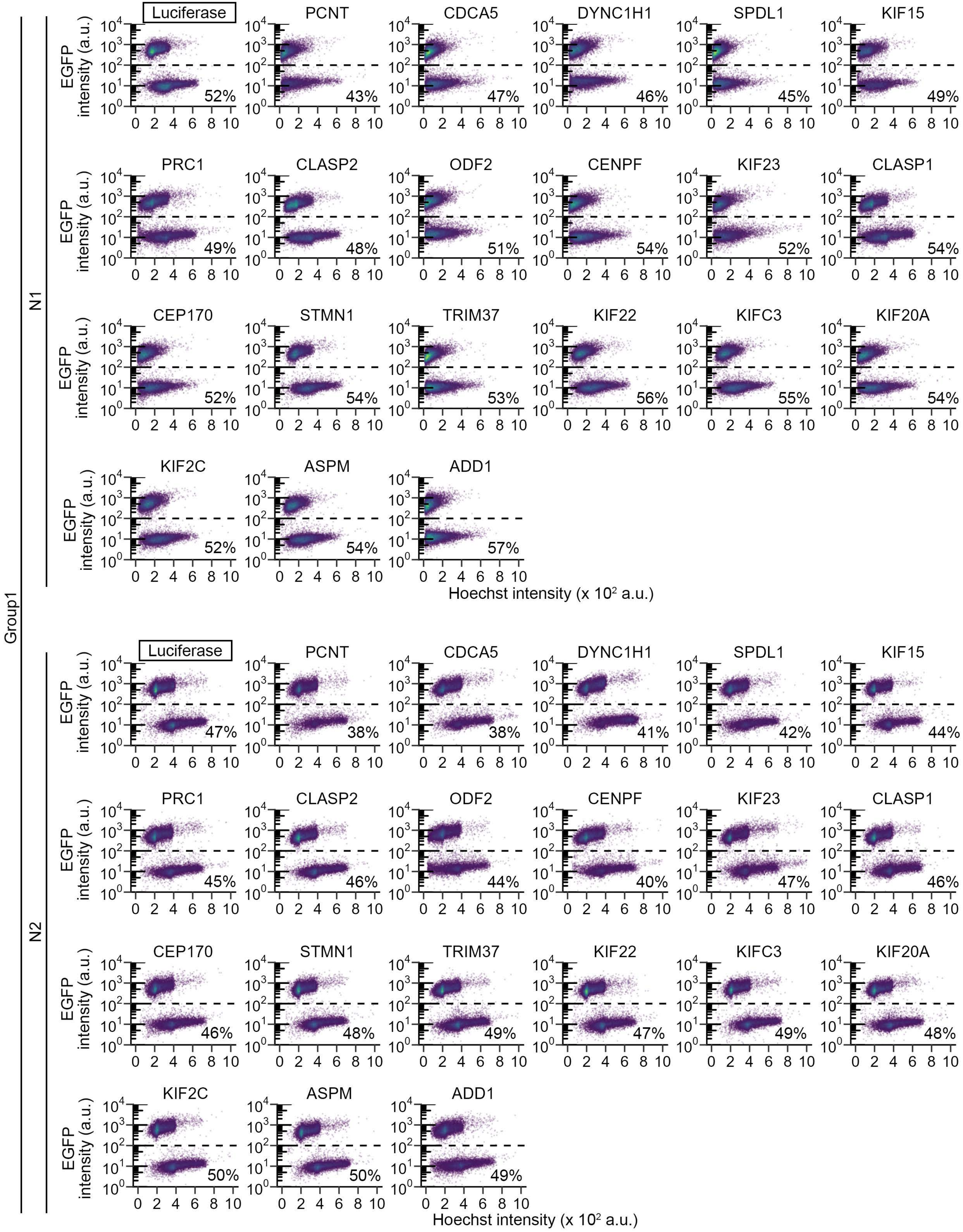


Supplementary materials page 1

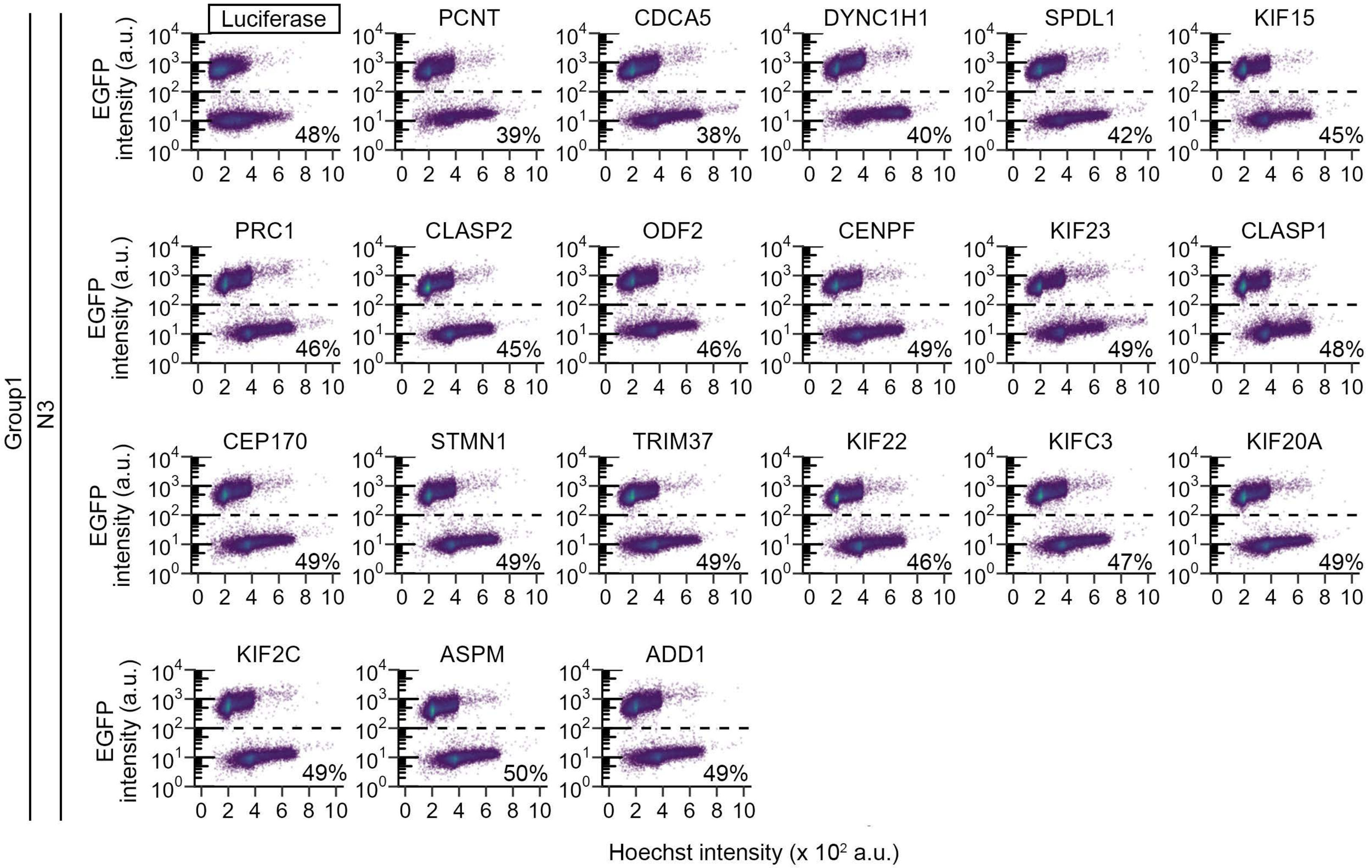

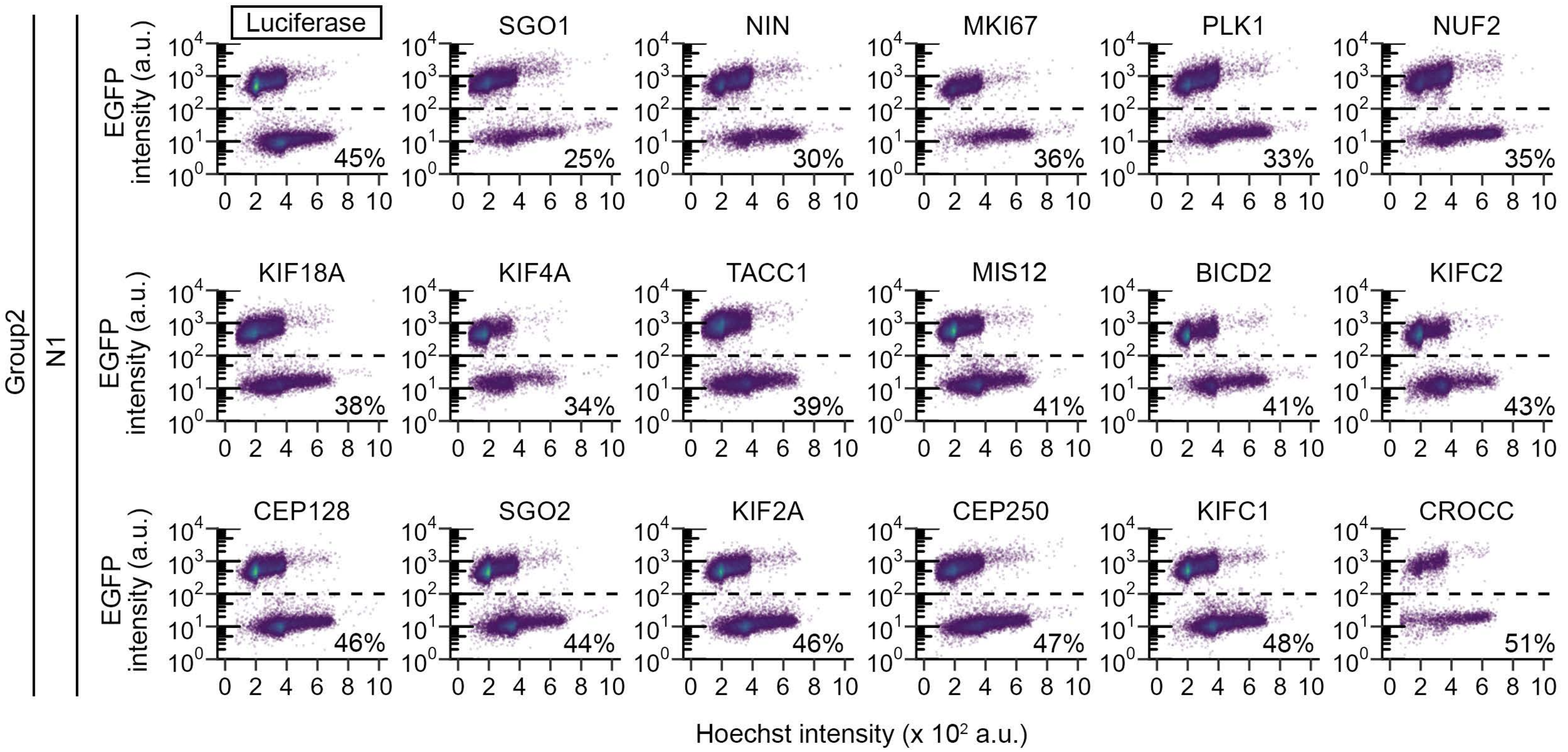

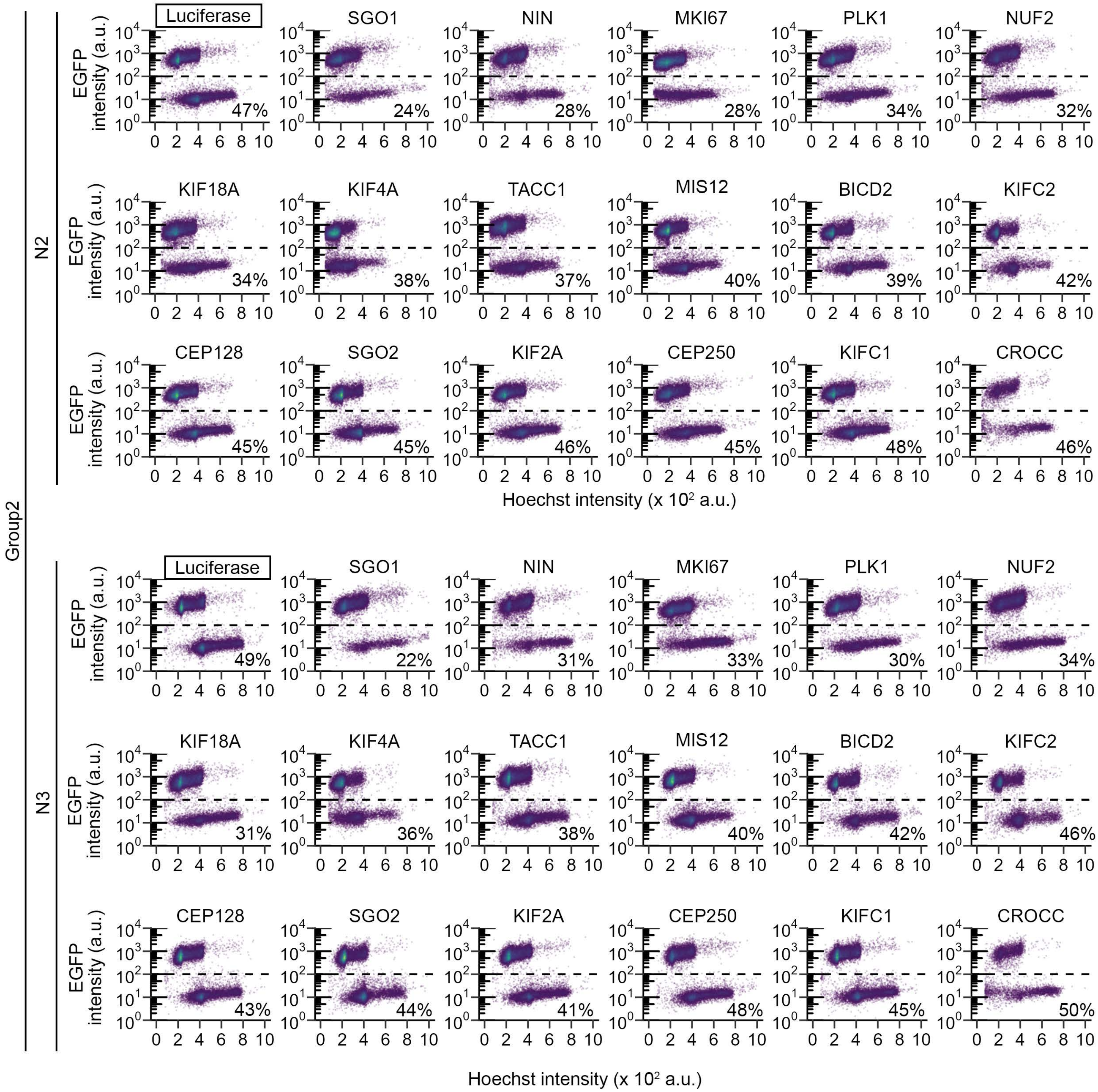

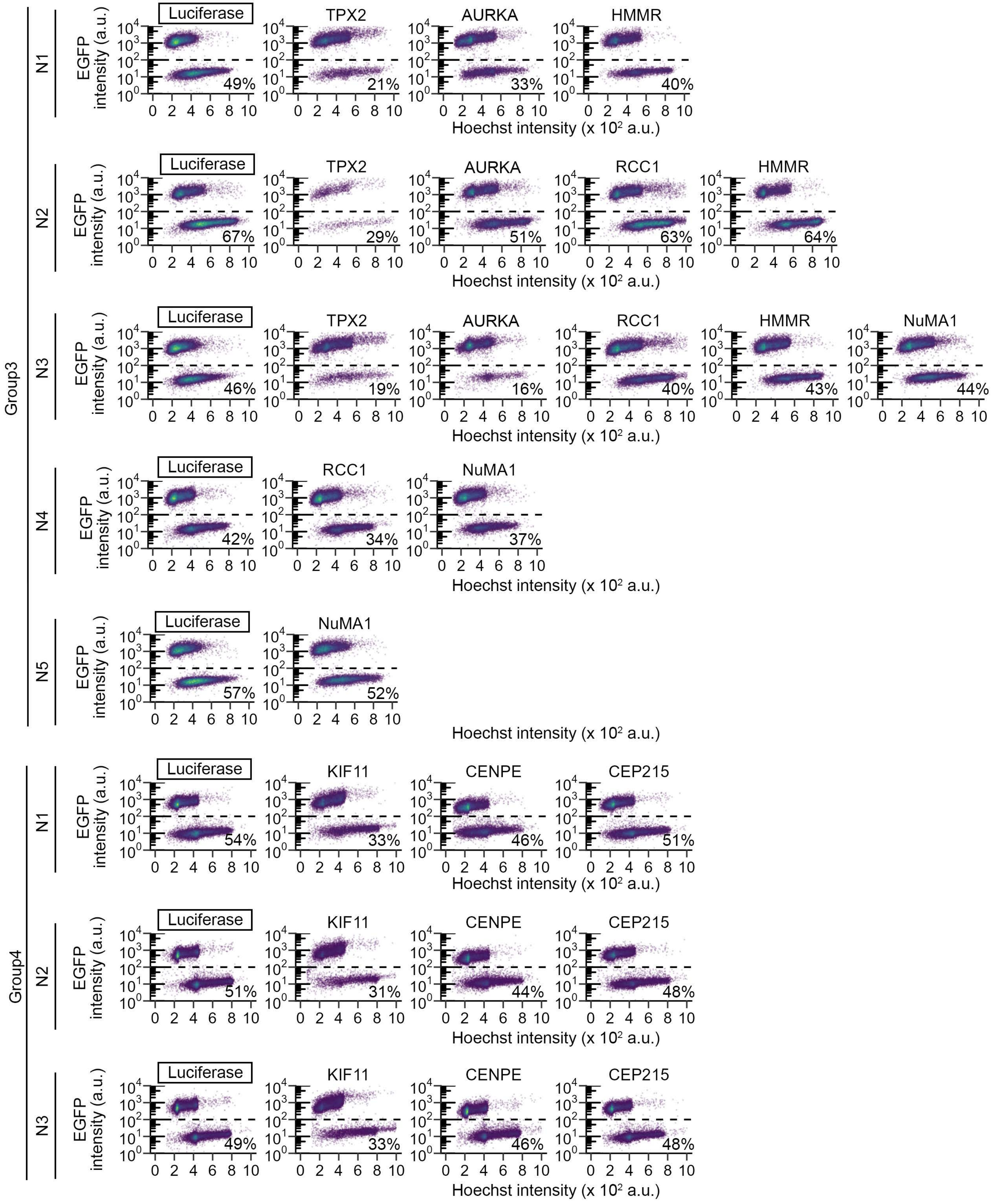

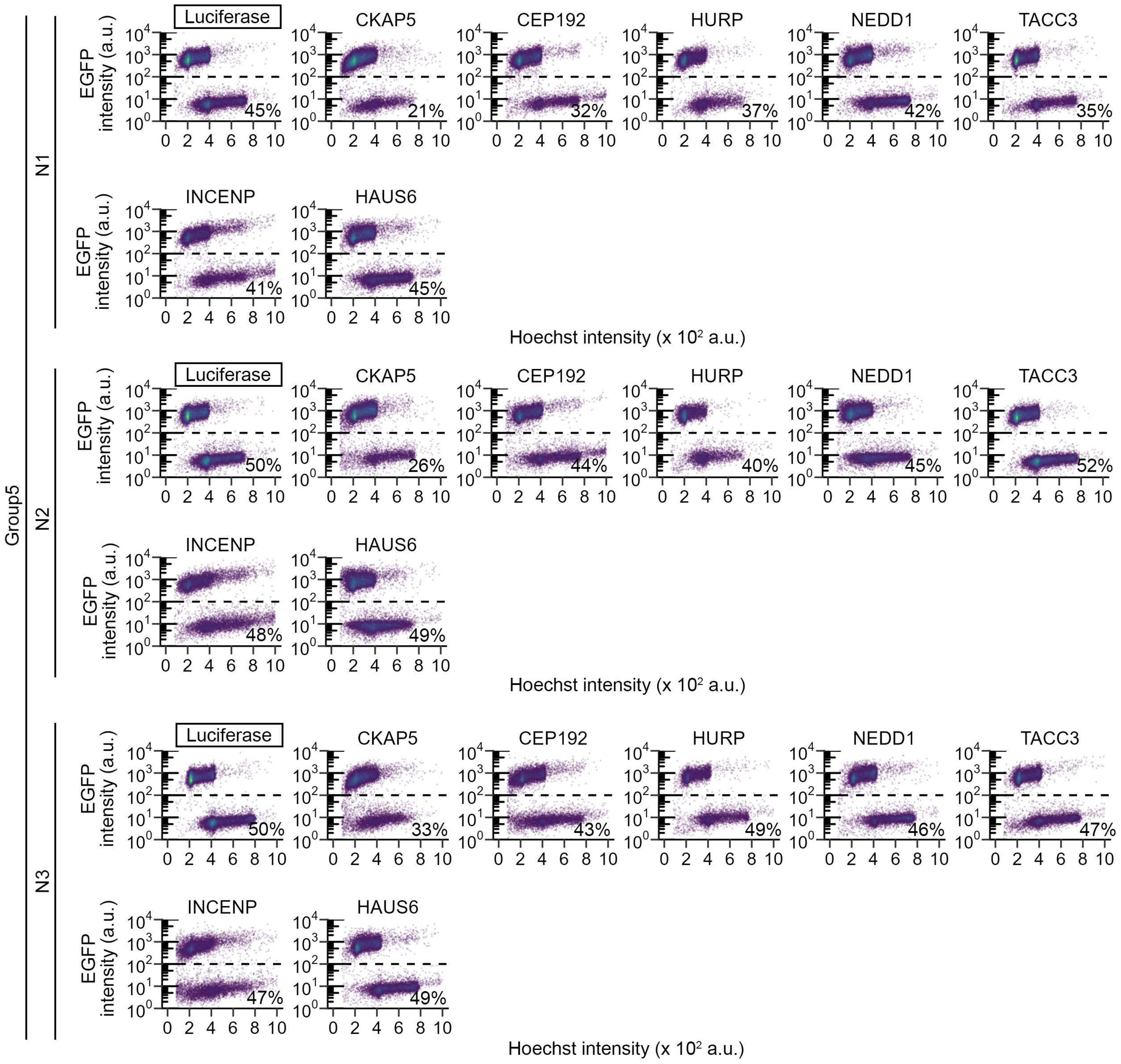


Supplementary materials page 5
