## Supplementary material S2 for "Spindle size scaling secures mitotic fidelity against whole-genome doubling-associated mechanical stress"

Immunoblotting associated with supplementary figure S1A

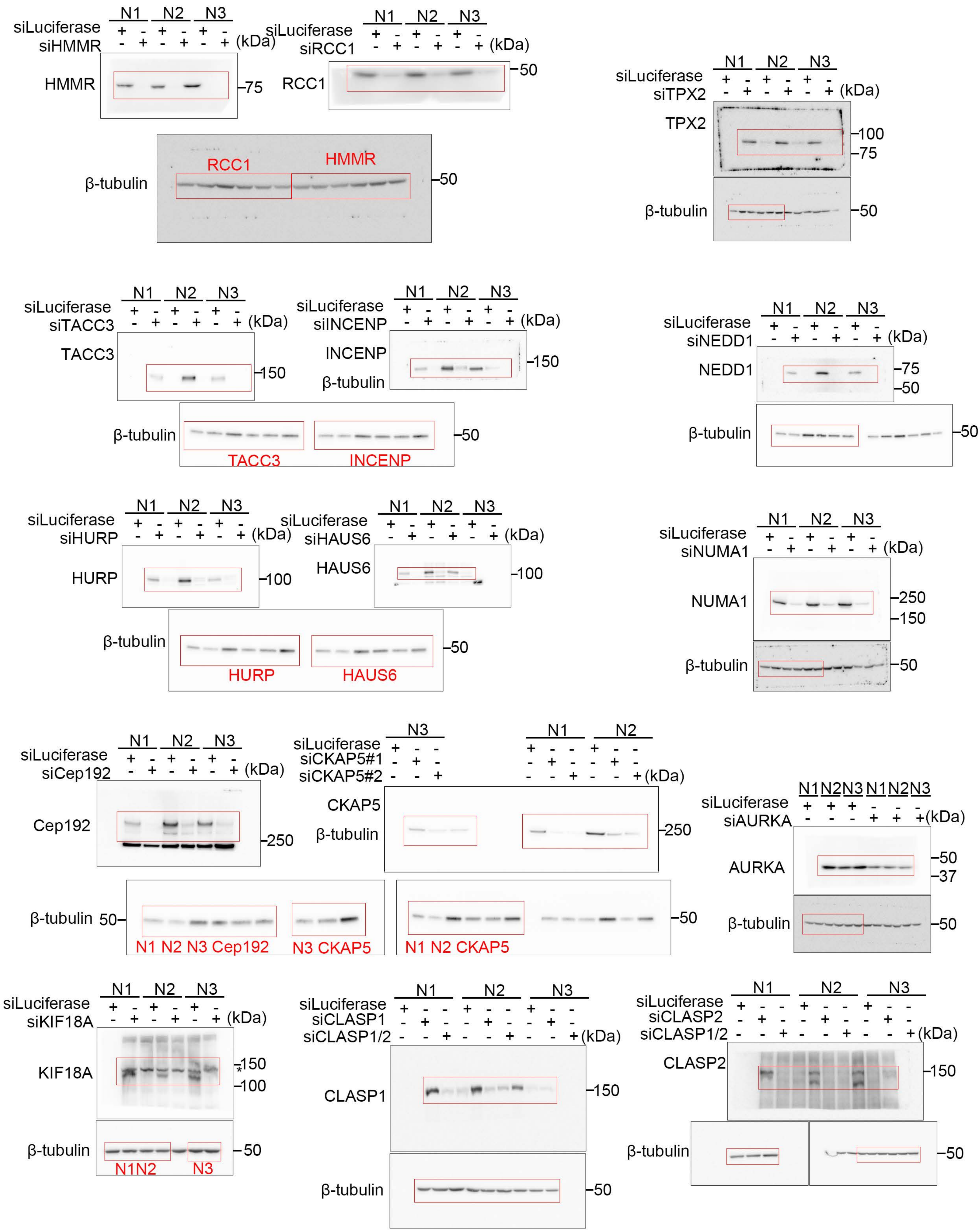

Immunoblotting associated with supplementary figure S2A

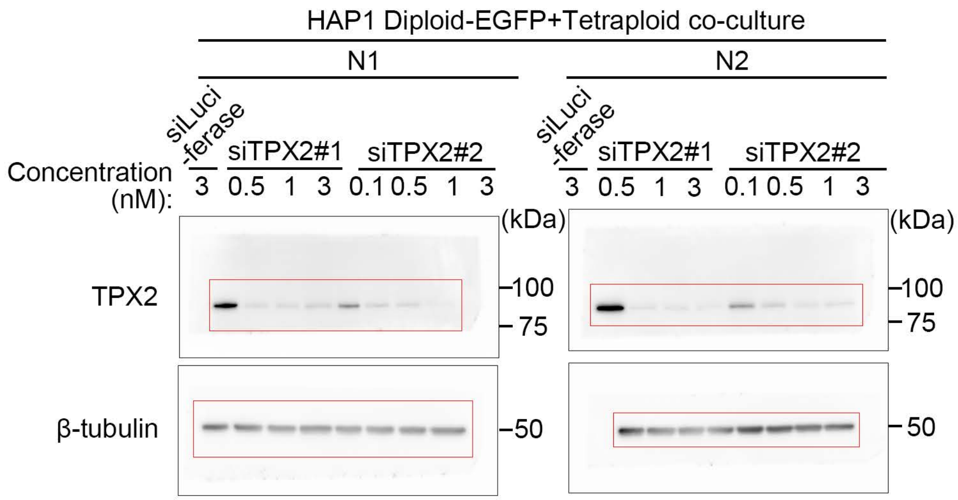


Immunoblotting associated with supplementary figure S2C

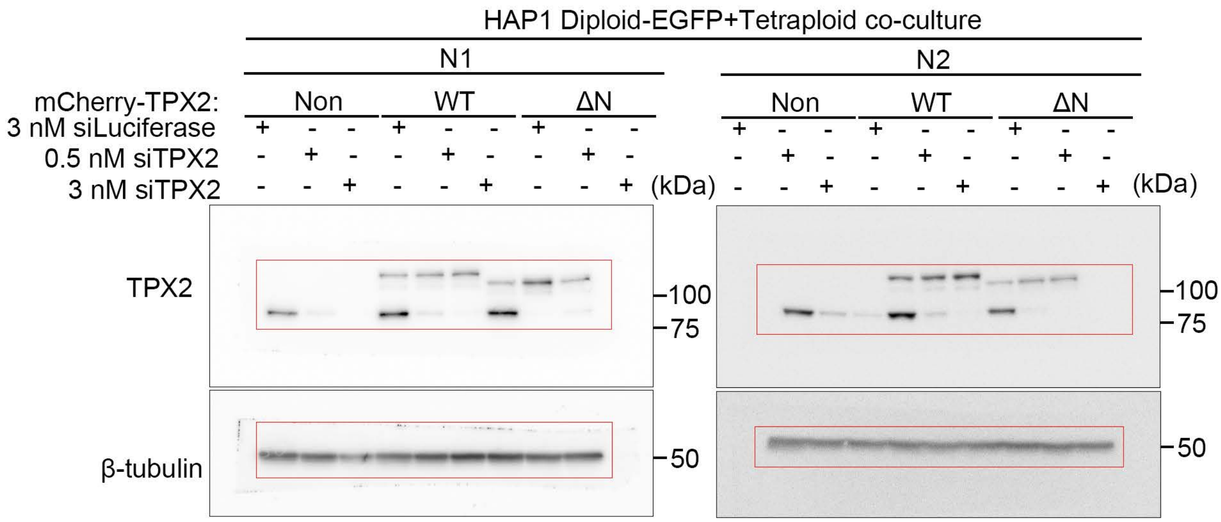

Immunoblotting associated with supplementary figure S3A

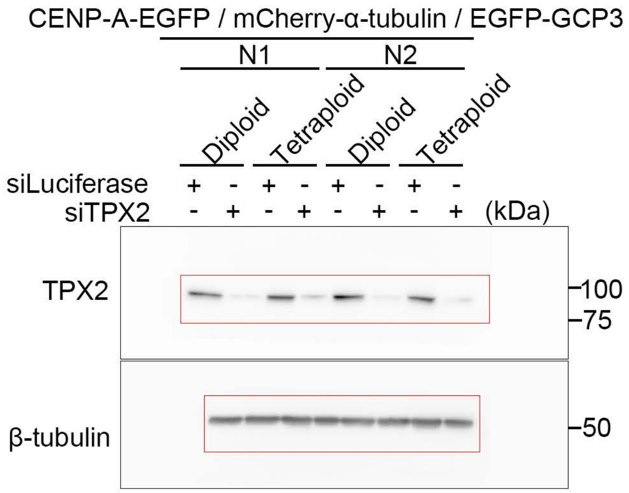


Immunoblotting associated with supplementary figure S3C

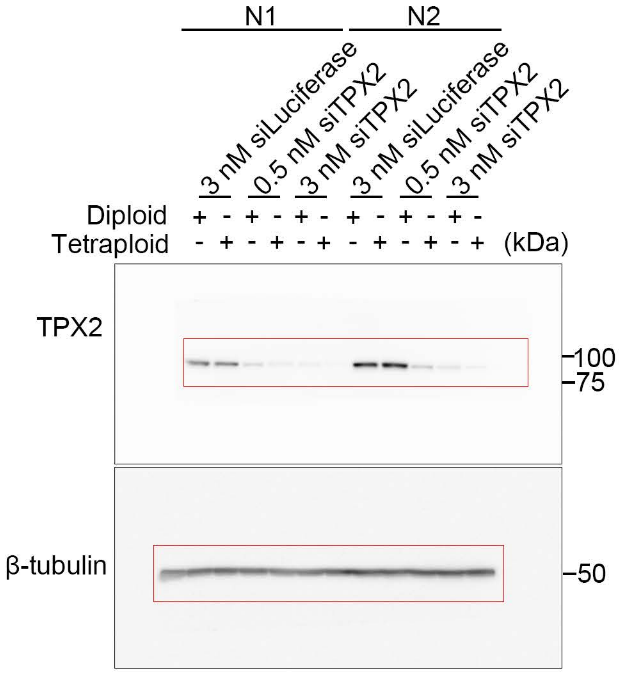


Immunoblotting associated with supplementary figure S3F

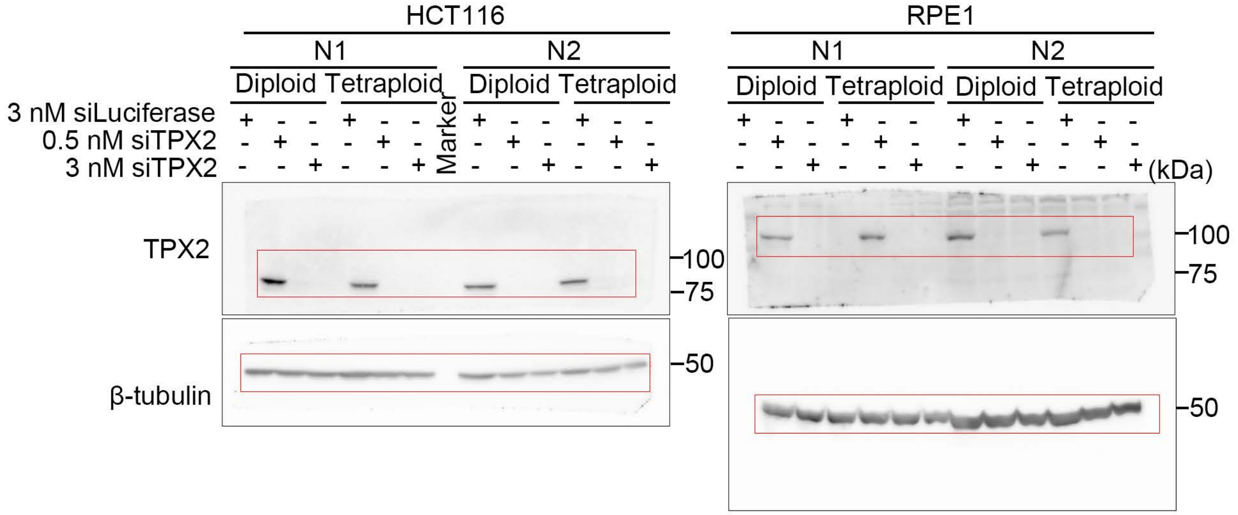

### Immunoblotting associated with supplementary figure S4C

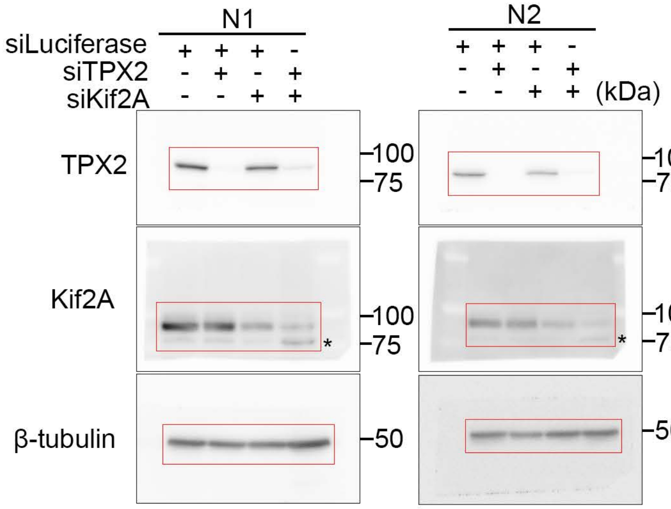


### Immunoblotting associated with supplementary figure S4E

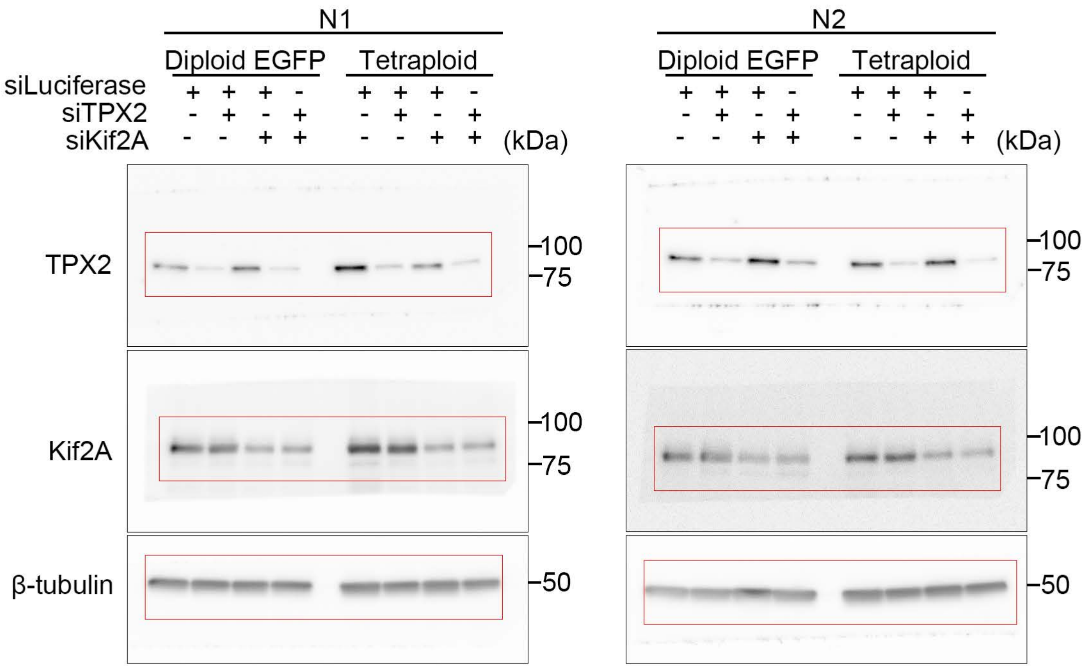

Immunoblotting associated with supplementary figure S5B (Top)

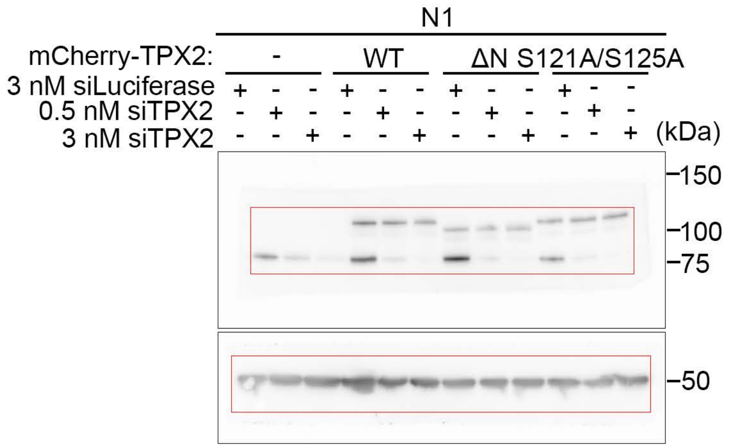


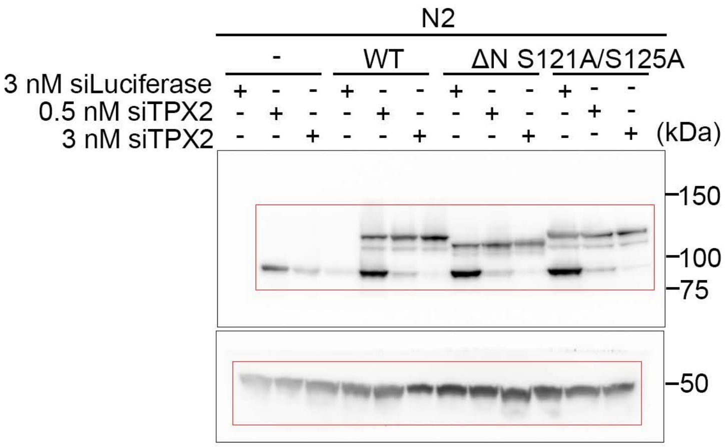


The N2 results are presented in supplementary figure S5B (Top)

Immunoblotting associated with supplementary figure S5B (Bottom)

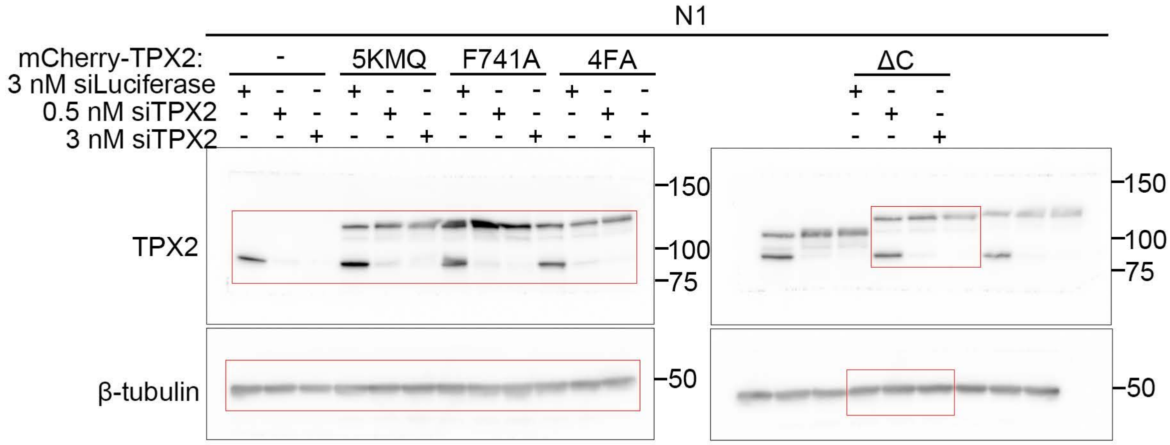


The N1 results are presented in supplementary figure S5B (Bottom)

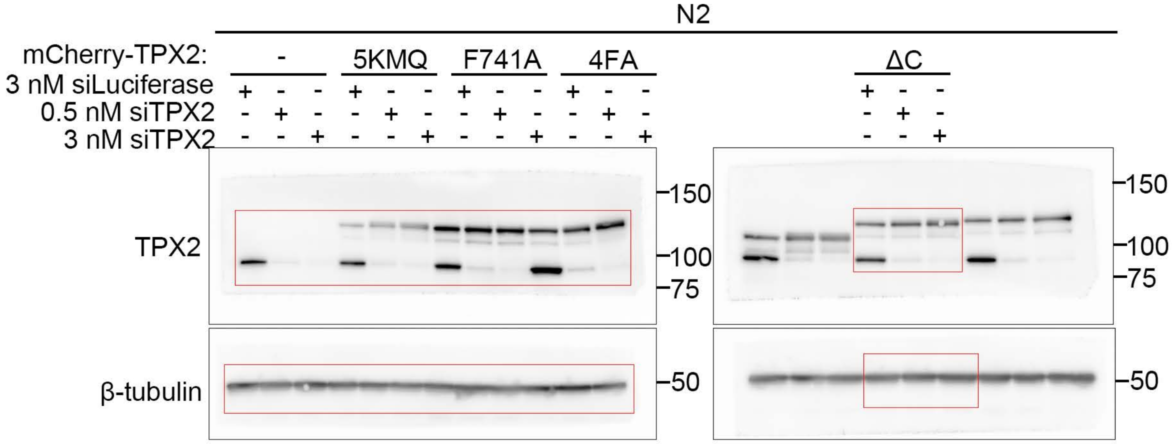

Immunoblotting associated with supplementary figure S5E

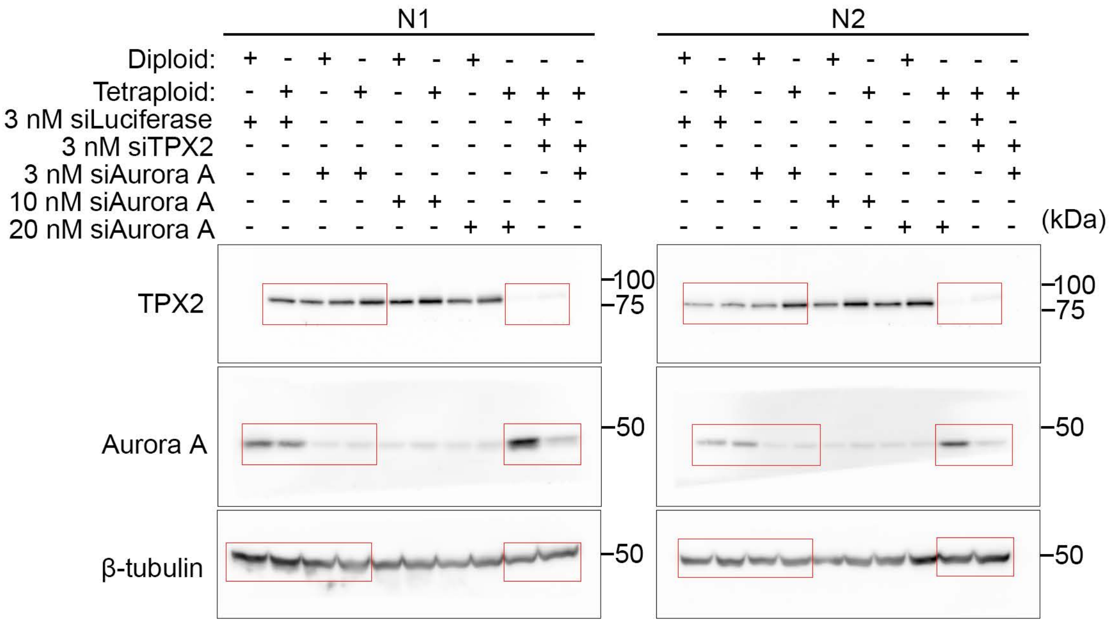

### Immunoblotting associated with supplementary figure S6A

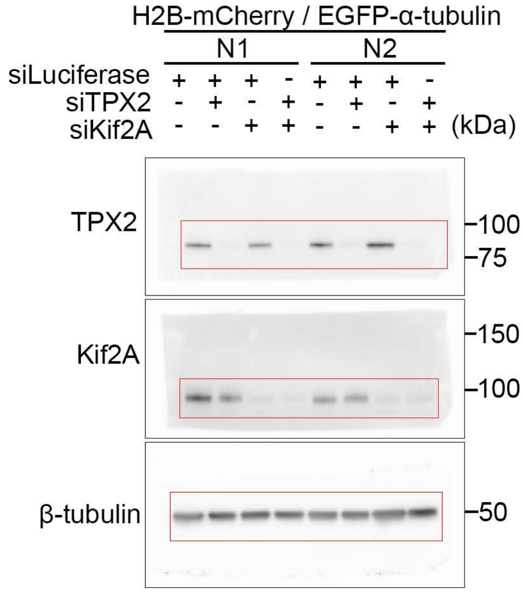


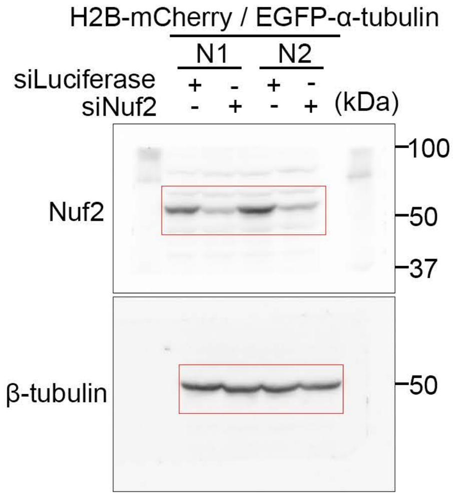


### Immunoblotting associated with supplementary figure S6H

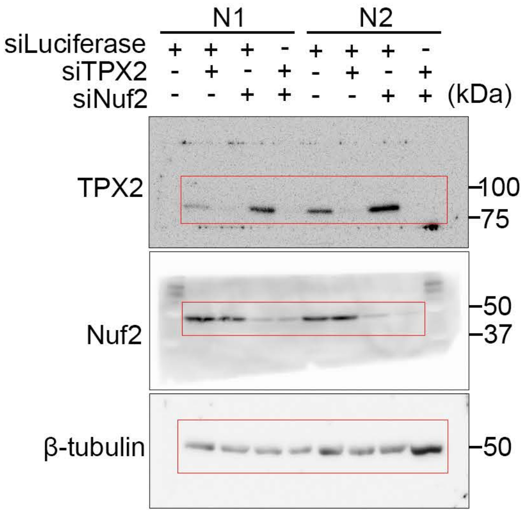


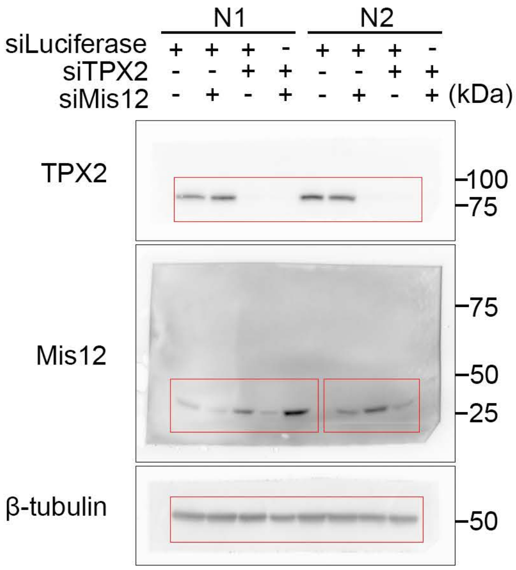

Immunoblotting associated with supplementary figure S7A

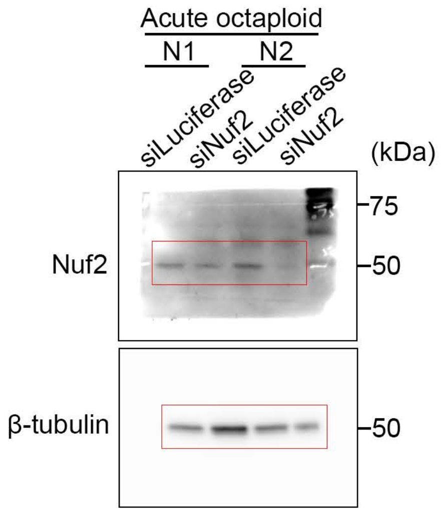


Immunoblotting associated with supplementary figure S7B

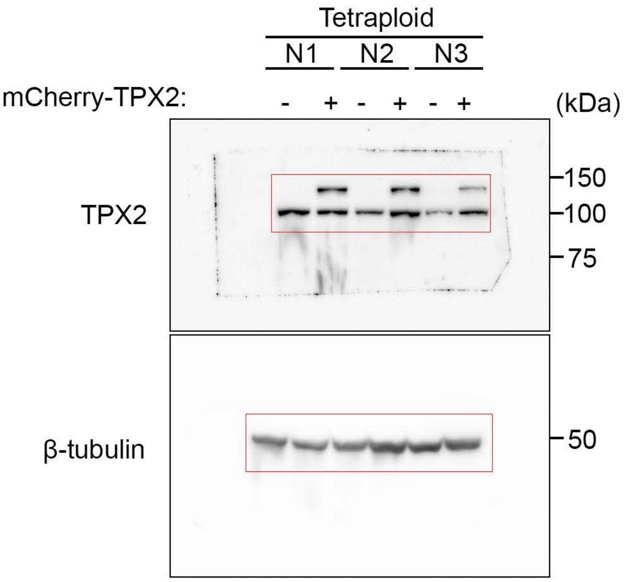
