## Supplementary Appendix Text for "Spindle size scaling secures mitotic fidelity against whole-genome doubling-associated mechanical stress"

**Supplementary Appendix S1**

We investigated the relationship between spindle structural features and recoiling dynamics after laser ablation under different ploidy levels and RNAi conditions. In addition to length and width, we quantified the global curvature of the contours of the outermost kinetochore fibers, or the angles between two outermost kinetochore fibers flanking equatorial chromosomes (θ_equator_) or a spindle pole (θ_pole_) (Fig. S6B). We quantified these parameters because these features may affect or reflect the spindle's mechanical state (*1, 2*).

Consistent with the results of the preceding live imaging analysis (Fig. 2), spindles were larger both in length and width in tetraploids compared to diploids (Fig. S6C). In tetraploids, θ_equator_ or θ_pole_ significantly decreased or increased, respectively, compared to diploids (Fig. S6C). Spindle contour curvature was significantly smaller in tetraploids than in diploids, possibly reflecting larger spindle sizes in tetraploids (Fig. S6C). TPX2 depletion and Kif2A depletion showed opposite effects on different structural features: Spindles in tetraploids became 69% in length and 84% in width by TPX2 depletion and 117% in length and 106% in width by Kif2A depletion (relative to the mock-depleted condition; Fig. S6D). TPX2 depletion significantly decreased θ_equator_, weakly increased θ_pole_, and significantly increased spindle contour curvature in tetraploids (Fig. S6D). In contrast, Kif2A depletion significantly increased θ_equator_, significantly decreased θ_pole_, and significantly decreased spindle contour curvature in tetraploids (Fig. S6D). Nuf2 depletion significantly increased spindle length without changing width, causing a substantial reduction in θ_pole_ in tetraploids (Fig. S6E).

We next analyzed the relationship among these structural features and their relationship to spindle recoiling dynamics across all experimental conditions (Fig. S6F). Spindle length showed correlations with θ_equator_ and θ_pole_: As spindle length increased, θ_equator_ or θ_pole_ tended to be larger or smaller, respectively (Fig. S6F). Among these structural features, spindle length showed the most evident (anti-) correlation with spindle recoiling velocity (Fig. S6F), supporting the idea that spindle elongation helps alleviate mechanical stress at spindle poles in tetraploids. On the other hand, changes in other structural features associated with spindle length complicate interpretation of these results. For example, reduced spindle contour curvature with spindle elongation may reduce potential bending stresses on outermost kinetochore fibers. A reduction in θ_pole_ may reduce the risk of pole splitting by weakening the component of tensile force acting on the pole in the spindle-width direction. However, because the forces within the spindle remain incompletely understood, we must consider the possibility that these geometric changes could affect polar stress through unidentified mechanisms. Future systematic analyses are needed to understand the mechanism by which polar mechanical stress decreases in correlation with spindle length.
